# PEATREST: A lifecycle assessment (LCA) model of carbon fluxes for restored afforested peatlands

**DOI:** 10.64898/2026.03.30.715261

**Authors:** J D. O’Sullivan, C Whittaker, G Xenakis, T Robson, M P. Perks

**Affiliations:** Forest Research, Bush Estate, Roslin, Midlothian, Scotland, UK, EH25 9SY

**Keywords:** Life cycle assessment, Carbon mitigation, Peatland restoration, Forest-to-bog restoration

## Abstract

Peatlands are an important terrestrial carbon sink which, when drained, can produce substantial CO_2_ efflux. Low productivity forestry planted on drained peatlands can become a net carbon source if losses from drained soils exceed sequestration by the trees. Decision support tools which assist resource allocation and intervention planning in forest-to-bog restoration are needed to mediate this substantial environmental harm. Predicting carbon mitigation benefits associated with forest-to-bog restoration is a major challenge, however, due to the lack of long-term monitoring programs and the fact that mitigation times depend on processes distant from the intervention. Here we introduce the PEATREST life cycle assessment (LCA) which predicts carbon fluxes associated with forest-to-bog restoration, including due to processes far from restored sites. The LCA estimates mitigation timescales defined as the time following intervention at which the restored peatland is predicted to sequester or store more carbon than the forestry would have if retained.

**Highlights:**

- Here we develop a novel forest-to-bog Life cycle assessment (LCA) tool
- The LCA predicts carbon mitigation times following peatland restoration
- The model combines a variety of process-based and empirical sub-models
- Example implementations for two different restoration scenarios are explored
- Sensitivity analysis highlights the model inputs that most impact outcomes

**Graphical abstract:** (A single, concise figure that serves as a visual summary of the main research findings described in your manuscript.)

The PEATREST Life cycle assessment (LCA) generates compound time series of carbon sequestration and carbon storage for two scenarios: the forest-to-bog peatland restoration (PR) and a counterfactual (CF) of forestry retention. By comparing the two scenarios, the LCA predicts the carbon mitigation timescales (vertical dashed lines). These are defined as the time following harvesting at which the peatland is predicted to sequester more (emit less), or to have stored more (lost less) carbon, than the forestry would have if retained.

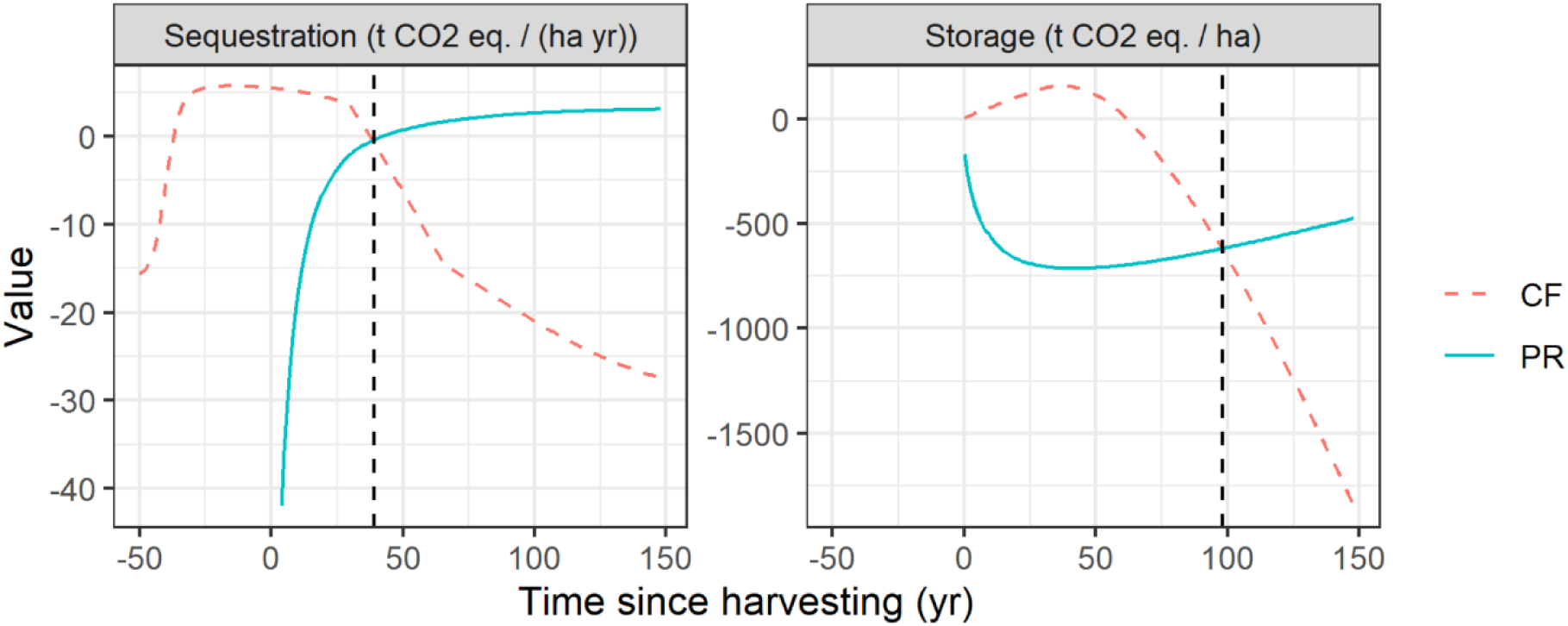

## 1. Introduction

Peatlands are an important terrestrial carbon store due to the slow decomposition of plant biomass within their anaerobic zones (Moore and Basiliko, 2006). When drained, for example for forestry, aerobic mineralisation of that stored carbon causes it to be rapidly lost to the atmosphere (Maljanen et al. 2010). At the global scale, an estimated 1Pg CO_2_ yr^-1^ is lost from drained peatlands (IPCC, 2014). In the UK, around 20% of the natural peatlands have been drained and replaced with conifer plantation (Anderson et al., 2016). Where interventions to restore peatland function raise the water table, peat forming (anaerobic) conditions can reduce emissions and, if successful in substantially restoring peatland function, can deliver long-term net greenhouse gas (GHG) mitigation (Mander et al. 2024). Peatlands are also broadly beneficial in biodiversity maintenance (Palmer, 2024), minimizing flood risk (Goudarzi et al. 2021), and ensuring safe drinking water (Xu et al. 2018).

There are a growing number of studies which give vital clues as to the typical timescales involved in forest-to-bog restoration. Water tables, for example, have been found to take >17 years (the maximum treatment length) to return to conditions typical of pristine bogs (Gaffney et al. 2018). Extrapolating from a time series of 24 years, the time to restoration of peatland vegetation was estimated to be in the range 50-100 years (Rydgren et al. 2025). Peatland status metrics have been shown to take 20-35 years to recover when enhanced restoration techniques are employed; longer when sites are allowed to recover with minimal intervention (Allan et al. 2024). Encouragingly, studies have shown the onset of zero net GHG flux and even net carbon sink dynamics 11-16 years after restoration (Hambley et al. 2018, Fundira et al. 2025) suggesting carbon mitigation benefits can occur prior to full recovery of ecosystem function. The lack of long-term, controlled experiments into peatland restoration dynamics is, of course, a major barrier to making empirically derived estimates of carbon mitigation times (Hughes et al. 2025), however work is on-going to fill that knowledge gap. Of particular note are studies of process-based soil carbon models (e.g. Smith et al. 2010) which may offer a computational route to assessing the lag times in restoration of ecosystem function that would otherwise take multiple decades to observe in experimental systems. Crucially, however, such studies are necessarily of narrow scope. An experimental assessment of carbon fluxes along a restoration gradient is unlikely to account for the lost sequestration potential of the forestry or the end use of the harvested woody biomass. To this end, the PEATREST life cycle assessment (LCA) tool was developed.

The goal of the PEATREST LCA is to construct two alternative time series of CO_2_ eq. emissions, one corresponding to the ‘counterfactual’ scenario in which the forestry is left in place, and the other summarising the various carbon fluxes, due to both management interventions and peatland ecosystem dynamics, associated with the forest-to-bog restoration. By comparing these time series, the model predicts two metrics representing the mitigation time: the carbon flux intercept *t*_*flux*_ and the carbon payback time *t*_*payback*_. The carbon flux intercept is the time following harvesting at which the rate of carbon sequestration by the peatland is first predicted to exceed that of the forestry, had it been left in place. The carbon payback time is the time following harvesting at which the peatland is predicted to have *stored* more carbon than the forestry would have if not removed.

## 2. Methods

### 2.1. Description of the LCA

The PEATREST LCA accounts for a variety of contributions to the net carbon flux for both the counterfactual and the restoration scenarios (Fig. 1). For the counterfactual, sequestration by the trees, emissions from the drained peats under the trees and leaching of aquatic carbon (DOC and POC) are represented. For the restoration scenario, the model accounts for one-off emissions associated with harvesting and restoration as well as emissions due to the decay of woody material which can either be left on site or removed and processed into wood products. Emissions rates from the peatland (including net negative emissions due to bog plant sequestration) are modelled along a restoration trajectory as are aquatic carbon losses. A full list of variables included in the PEATREST LCA is given in Appendix A.

**Fig. 1.**
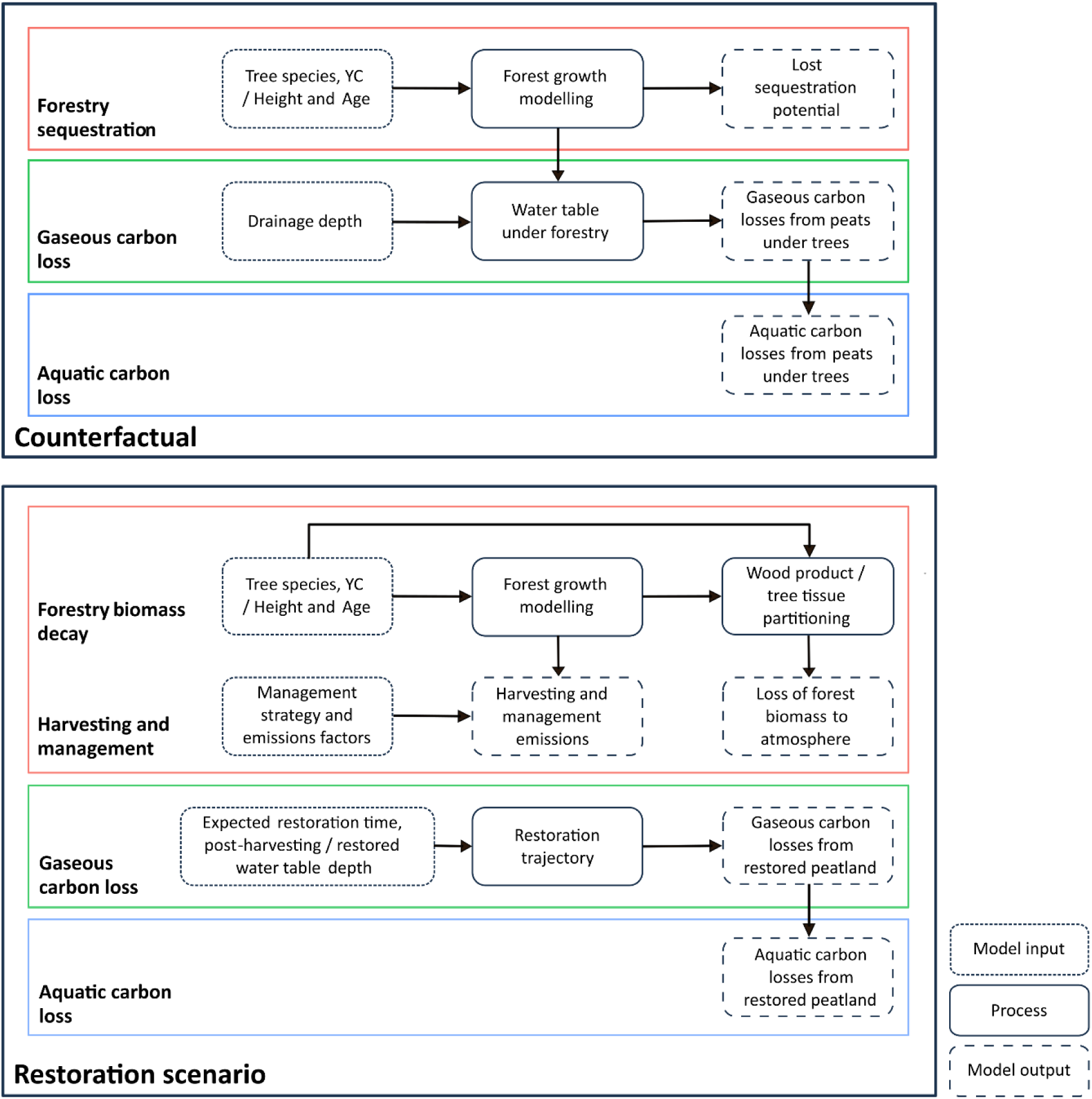
Schematic representation of the PEATREST LCA.

The LCA makes extensive use of two models commonly used in applied forestry: 3PG (Landsberg and Waring, 1997) and CARBINE (Thompson and Matthews, 1989). 3PG is a forest growth model which is used to predict carbon sequestration by the forestry. This sequestration defines both the lost sequestration potential but also the total carbon stored in the forestry at the time of harvesting. CARBINE is a carbon accounting model which predicts the amount of harvested forestry biomass allocated to various wood products (biofuels, paper, panelling and sawn wood) or tissue types (root, stem, branch and foliage) as a function of stand age and yield class. These are used to model the carbon stocks at the time of harvesting. These carbon stocks then represent the initial state for a set of decay functions representing the loss of forest carbon to the atmosphere over time.

### 2.2. Emissions rates modelling

For both the counterfactual and the restoration scenarios, it is necessary to estimate peatland emissions rates including sequestration (i.e. negative emissions). CO_2_ and CH_4_ emissions rates from peatlands as a function of water table depth are modelled using the following empirical equations (Evans et al. 2021):

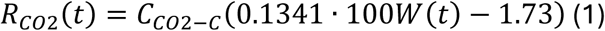

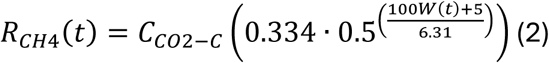

*R*_*CO*2_(*t*) and *R*_*CH*4_(*t*) represent emissions rates (t CO_2_ eq. ha^-1^ yr^-1^), and *W*(*t*) (m) is the annual average water table depth at time *t*. *C*_*CO*2−*C*_ = 3.667 converts losses of carbon bound in CO_2_ molecules to CO_2_ equivalent masses. The multiplication by 100 is required to convert from m to cm. Emissions rates are converted into total emissions *L*_*tot*_(*t*) = *L*_*CO*2_(*t*) + *L*_*CH*4_(*t*) (t CO_2_ eq.) by multiplying by the area harvested at the timestep length (1 yr).

### 2.3. Forest growth modelling

Sequestration from the forestry is represented using a simplified version of the 3PG forest growth model (Landsberg and Waring, 1997; Smith et al., 2011). The photosynthetically available light absorbed by the forestry (MJ m^-2^ yr^-1^) is modelled as

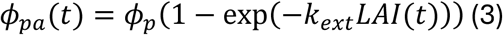

Here *ϕ*_*p*_(MJ m^-2^ yr^-1^) is the average annual incident radiation above the canopy which is fixed at 2880 MJ m^-2^ yr^-1^ based on UK annual average solar irradiance. *k*_*ext*_is the unitless light extinction coefficient, and *LAI*(*t*) is the unitless leaf area index of the forestry. *LAI*(*t*) is a dynamic variable given by

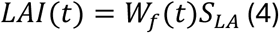

*W*_*f*_(*t*) is the foliage biomass at time *t* (t C ha^-1^), while *S*_*LA*_is the specific leaf area of the focal species (m^2^ kg C^-1^). The foliage biomass dynamics are given by the difference equation

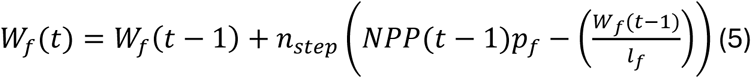

*NPP*(*t* − 1) (t C ha^-1^) is the net primary productivity for the previous timestep, *p*_*f*_is the proportional carbon allocation to the leaves, and *l*_*f*_ the leaf longevity. *n*_*step*_is the timestep length, set to 1 year.

Net primary productivity *NPP*(*t*) is given from the gross primary productivity *GPP*(*t*) (t C ha^-1^) scaled by fixed ratio *Y* = 0.47 (Waring et al. 1998).

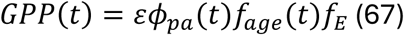

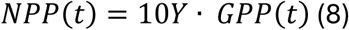

The multiplication by 10 converts from units of kg m^-2^ to t ha^-1^. *ε* is the maximum light use efficiency for the tree species (kg C MJ^-1^), while *f*_*age*_(*t*) and *f*_*E*_are the unitless age and environmental growth modifiers respectively. The age modifier, which ensures sequestration rates drop off for old stands, is given by

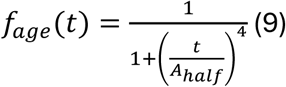

Here, *t* is the stand age (yr) while *A*_ℎ*alf*_is the age (yr) at which there is a 50% decrease in light use efficiency (i.e. *f*_*age*_(*t*) = 0.5). Unlike ‘full’ 3PG simulation models, this simplified scheme does not incorporate explicit, time dependent environmental inputs. Instead, the constant environmental growth modifier is set so as to reproduce the sequestration rates corresponding to specific yield classes (Fig. 2, see Appendix B for details on setting *f*_*E*_). A full list of model parameters for the simplified 3PG model is given in Tab. 1.

**Fig. 2.**
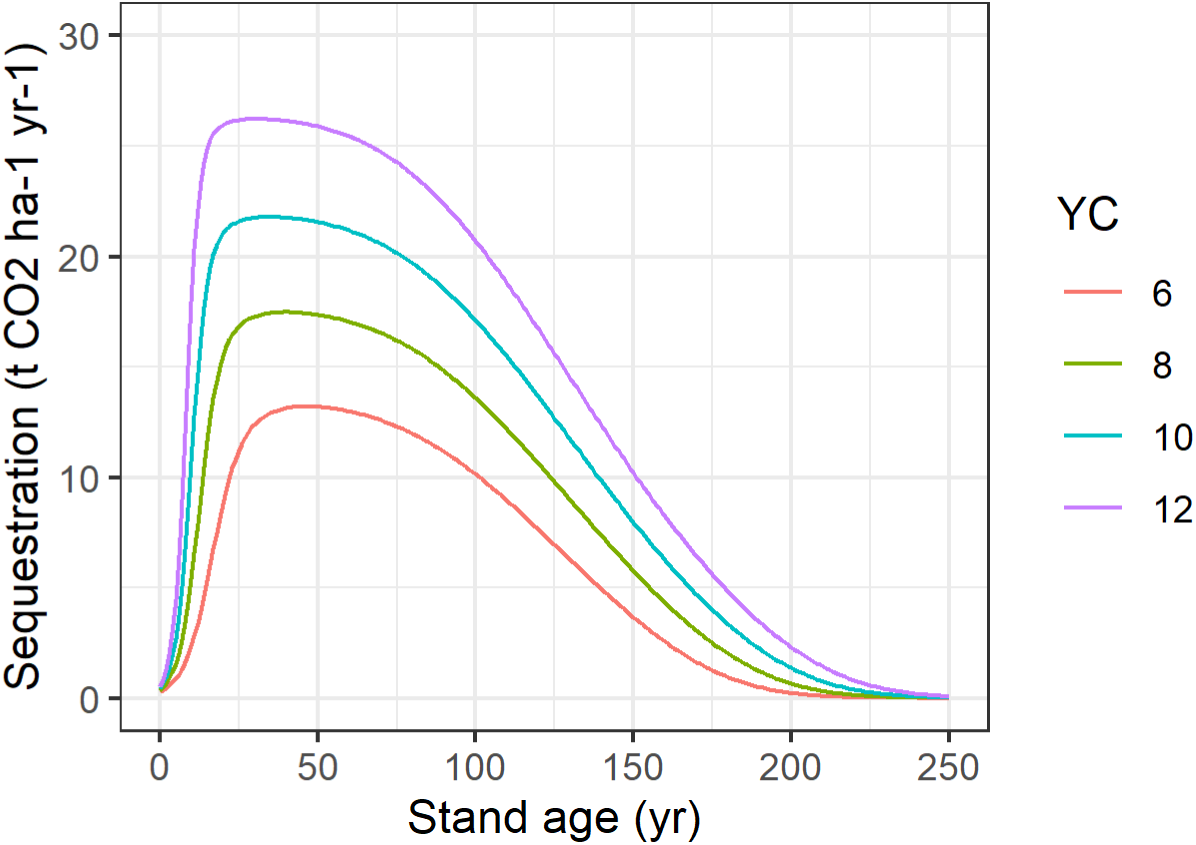
Sequestration rates (t CO_2_ ha^-1^ yr^-1^) predicted using the simplified 3PG model parameterised to represent Sitka spruce stands of yield class 6-12.

**Tab. 1.** Parameters used in the simplified 3PG model, adapted from Smith et al. (2011).

| Parameter | Scots pine | Sitka spruce | Units |
| --- | --- | --- | --- |
| Average annual photosynthetically active radiation, $\phi_p$ | 2880 | | MJ m<sup>-2</sup> yr<sup>-1</sup> |
| Light extinction coefficient, $k_{ext}$ | 0.5 | | - |
| Specific leaf area, $S_{LA}$ | 6 | 8 | m<sup>2</sup> kg C<sup>-1</sup> |
| Leaf longevity, $l_f$ | 4 | 7 | yr |
| Maximum light-use efficiency, $\varepsilon$ | 0.00152 | 0.001656 | kg C MJ<sup>-1</sup> |
| Ratio of NPP to GPP, $Y$ | 0.47 | | - |
| Proportional carbon allocation to foliage, $p_f$ | 0.25 | | - |
| Age for 50% reduction in light-use efficiency, $A_{half}$ | 100 | 120 | yr |

### 2.4. Emissions from soils under trees

The counterfactual scenario must consider emissions from the drained peats on which the forestry is planted, *L*_*forest*_(*t*). For low yield class forestry, these emissions may exceed sequestration rates in the trees, thus making the forestry a net carbon source (Anderson, 2020). The same equations used for estimating emissions rates from undeveloped or restored peatlands (Eqs. (1) – (2)) are used to predict emissions from drained peats under the trees. This makes the simplifying assumption that the hydrological effect of the trees on the peats dominates any alternative impacts they may have on emissions rates (e.g. via nutrient input, substrate compaction, etc.).

To estimate emissions from peats under trees, water table depths under the forestry need to be modelled and projected forward beyond the point of harvesting. These are estimated by assuming rooting depth serves as a reasonable proxy for water table depth. Rooting depth as a function of yield class and stand age is given from the root biomass estimated by CARBINE. Initial estimates of rooting depths *d*_*r*_^′^(m) are given by the equation

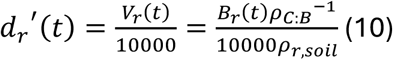

*V*_*r*_(*t*) is the volume of the rooting zone (m^3^), *B*_*r*_(*t*) (t Biomass ha^-1^) is the root biomass and *ρ*_*r*,*soil*_ (t Biomass m^-3^ Soil) is the biomass density of the rooting zone (tonnes root biomass in 1 m^3^ soil). Dividing by 10000 (m^2^) converts the volume (soil surface area of 1 ha) to an average depth (m). Based on analysis of tree pulling data from peatlands (Nicoll et al. 2006), *ρ*_*r*,*soil*_ is set to 0.060 (0.390 – 0.080) t Biomass m^-3^ Soil. *ρ*_*C*:*B*_ is the unitless Carbon: Biomass ratio of the forestry, set to 0.5 t C t^-1^ Biomass by default (Matthews, 1993). This is required since CARBINE predicts biomasses in oven dried tonnes, whereas the tree pulling data records root plate mass in green biomass units.

Initial water table depths are estimated from the user input drainage depth *d*_*d*_. Final water table depths are set by the smaller of *d*_*r*,*max*_, the maximum rooting depth, or *d*_*p*_, the peat depth of the site, i.e. min(*d*_*p*_, *d*_*r*,*max*_). Additional increases in water table depth beyond *d*_*p*_are possible but are not expected to impact emissions rates. Thus, the water table depth under the trees as a function of stand age is modelled using a bounded linear equation of the form

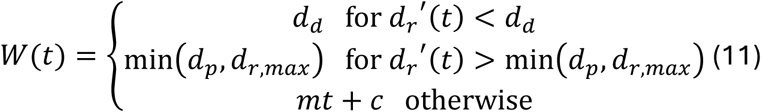

*d*_*r*,*max*_was also estimated from tree pulling data (Nicoll et al. 2006), in which no significant association between yield class and rooting depth was found, and set to 0.618 (0.390 – 0.845) m. This value corresponds to the depth of the root plate, since fine roots are much harder to measure. The impact of fine roots on the hydrology of the soil unavoidably contributes to uncertainty in the model. The coefficients *m* and *c* for a given yield class are estimated by simple linear regression. An example of the procedure is shown in Fig. 3. Note that in reality, additional increases in root biomass once max root depths are attained are likely to correspond to increases in root density and the width of the root plate. This effect is represented implicitly via the bounded linear root depth model.

**Fig. 3.**
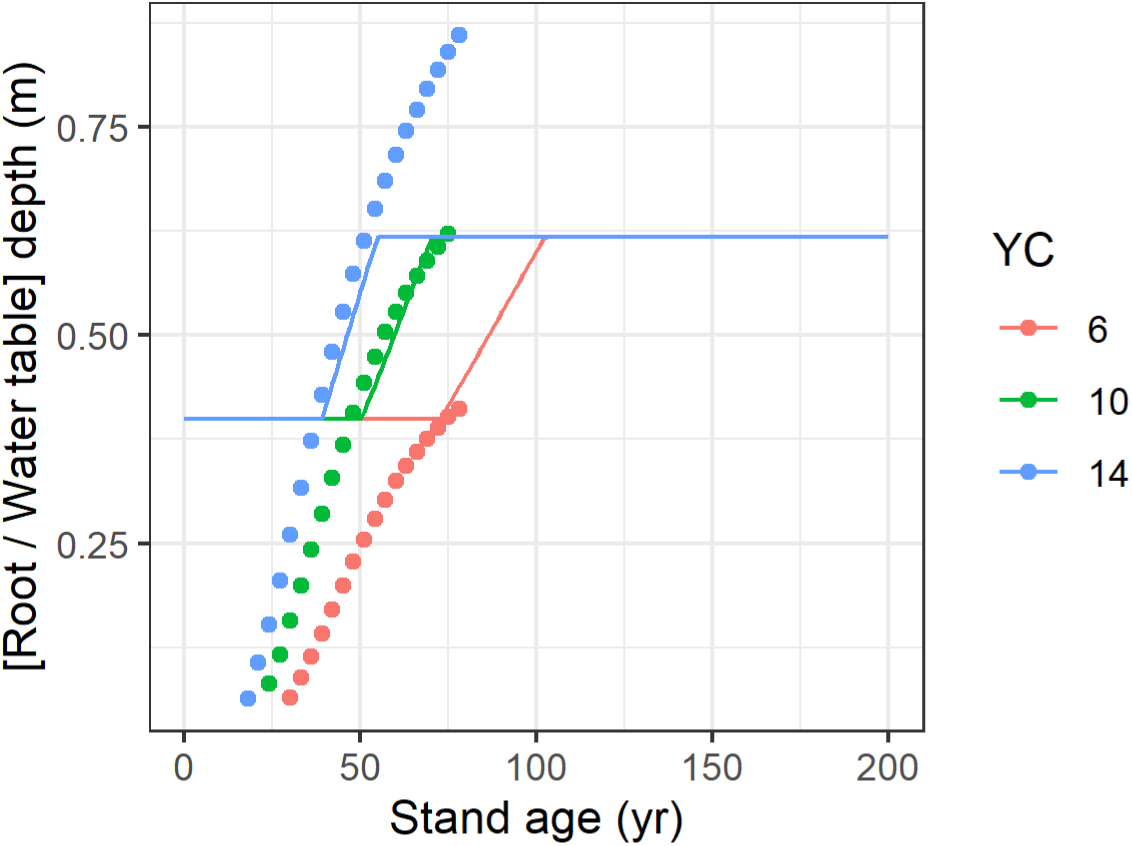
Root / water table depths for various yield classes estimated from a combination of CARBINE predictions of below ground biomass (Eq. (10), points) and bounded linear regression (Eq. (11), lines). Initial water table depths due to peat drainage set to 0.4 m (as in Hermans et al. 2022). Max root depths and biomass densities of the rooting zone set to average values (0.618 m and 0.06 t Biomass m^-3^ Soil respectively, Nicoll et al. 2006) in all cases since no significant association was found with yield class.

### 2.5. Aquatic carbon losses (DOC/POC)

There is evidence that DOC flux rates for undeveloped and afforested peats are broadly consistent (Buckingham et al. 2008; Melniks et al. 2024). This suggests the same model for aquatic carbon losses can be applied to both the afforested and restored scenarios. Previous carbon accounting studies have represented DOC and POC losses from drained peats by simple ratios to the total gaseous emissions (in t CO_2_ eq.). Smith et al. (2011) use empirically estimated ratios of *ρ*_*DOC*_ = 0.26 (0.07 – 0.40) for DOC fluxes and *ρ*_*POC*_ = 0.08 (0.04 – 0.10) for POC fluxes. These give a total aquatic carbon leaching rates of *ρ*_*AqC*_ = 0.34 (0.11 – 0.50) times the total gaseous losses from the peats. This approach cannot be used for the case that net peatland emissions are negative e.g. for sites predicted to become net carbon sinks following restoration. Therefore, a hybrid approach has been applied in the PEATREST LCA. Aquatic carbon losses to atmosphere *L*_*AqC*_ (t CO_2_ eq. ha^-1^ yr^-1^) are represented using a bounded linear function parameterised as in Smith et al. (2011), but capped at empirically observed rates:

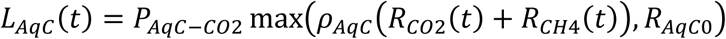

*P*_*AqC*−*CO*2_is the unitless proportion of aquatic carbon that ultimately ends up as atmospheric CO_2_. This is set to a value of 0.6 (0.4 – 0.8) (IPCC, 2001) with the remaining carbon assumed fixed in ocean floor sediments. *R*_*AqC*0_ is a measured average aquatic carbon loss from British peatlands, set to 0.338 (0.041 – 0.635) t CO_2_ eq. ha^-1^ yr^-1^ (average values of 0.281 and 0.058 t CO_2_ eq. ha^-1^ yr^-1^ for DOC and POC fluxes respectively, Worrall et al., 2009; Dawson et al., 1995; Dawson et al., 2002).

### 2.6. Silvicultural/restoration emissions

#### 2.6.1. Emissions due to management activities

The model accounts for two types of emissions due to silvicultural and restoration practices. Discrete, one-off emissions due to mechanised interventions are considered, as are continuous-time emissions due to the decay of either woody products (off-site timber utilisation) or forest biomass left to decompose on site. Discrete emissions associated with management of the site are represented using empirically estimated emissions factors (Tab. 2). Emissions due to a given intervention are estimated by multiplying emissions factors by one or more of: i) total area harvested/restored (user input); ii) total above ground tree biomass/volume (predicted using CARBINE); iii) distance from the site to the corresponding processing location (user input). The LCA includes the option to model a fallow period between harvesting and restoration. If this is the case, restoration emissions occur at time *t*_*fallow*_(yr) after harvesting. Further details including the sources of the estimates shown in Tab. 2 are given in Appendix C.

**Tab. 2.** Emissions factors for the various mechanised interventions available in the LCA. Note that mulching tractors operate on standing trees therefore if mulching is selected, harvesting emissions are omitted.

| Intervention | Symbol | Emissions factor |
| --- | --- | --- |
| Transportation | $E_{transport}$ | 39.33 (38.50 – 40.15) g CO<sub>2</sub> km<sup>-1</sup> t<sup>-1</sup> |
| Harvesting | $E_{harv}$ | 3.32 t CO<sub>2</sub> m<sup>-3</sup> |
| Mulching | $E_{mulch}$ | 302.8 (607.0 - 911.2) kg CO<sub>2</sub> ha<sup>-1</sup> |
| Bund creation | $E_{bund}$ | 25.5 (18.75 – 75.0) kg CO<sub>2</sub> ha<sup>-1</sup> |
| Drainage damming | $E_{dam}$ | 0.91 (0.46 – 1.86) kg CO<sub>2</sub> ha<sup>-1</sup> |
| Furrow smoothing | $E_{smooth}$ | 385.25 (373.32 – 397.18) kg CO<sub>2</sub> ha<sup>-1</sup> |
| Turfing (locally sourced) | $E_{turf,local}$ | 417.46 kg CO<sub>2</sub> ha<sup>-1</sup> |
| Turfing (imported) | $E_{turf,import}$ | 1516.42 kg CO<sub>2</sub> ha<sup>-1</sup> |
| Fertilising | $E_{fert}$ | 1983.37 kg CO<sub>2</sub> ha<sup>-1</sup> |

#### 2.6.2. Emissions from wood products and in situ decomposition

Emissions rates associated with the decay of forest biomass depend on whether the material is removed from the site and processed into wood products (paper, panel or timber) or left to decompose on site. In all cases, an exponential decay function is used to represent the losses of carbon bound up in forest biomass to the atmosphere. The loss to the atmosphere at time *t* from wood product or tissue type *X*, *L*_*X*_(*t*) (t CO_2_) is modelled as the difference between the carbon currently stored in that product *S*_*X*_(*t*) (t CO_2_ eq.) and the corresponding quantity from the previous timestep *S*_*X*_(*t* − 1):

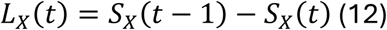

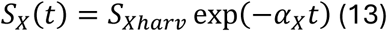

The decay rates *ɑ*_*X*_ are given from the half-life *λ*_*X*_ (estimated from literature) of the corresponding wood product or tissue type by

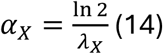

For processed material, the initial *decomposable* carbon pool stored in each product *S*_*X*ℎ*arv*_ is given by

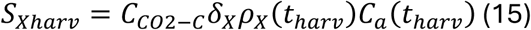

*C*_*a*_(*t*_ℎ*arv*_) is the above-ground woody carbon stored by the trees at the time of harvesting and *ρ*_*X*_(*t*_ℎ*arv*_) is a unitless ratio estimated from CARBINE, which gives the proportion of above-ground biomass allocated to product *X* if harvested at time *t*_ℎ*arv*_for a given species/yield class. Note that for larger yield classes, the proportion of above ground biomass allocated to slow decaying wood products is higher, while the proportion allocated to biofuels is lower (Fig. 4).

**Fig. 4.**
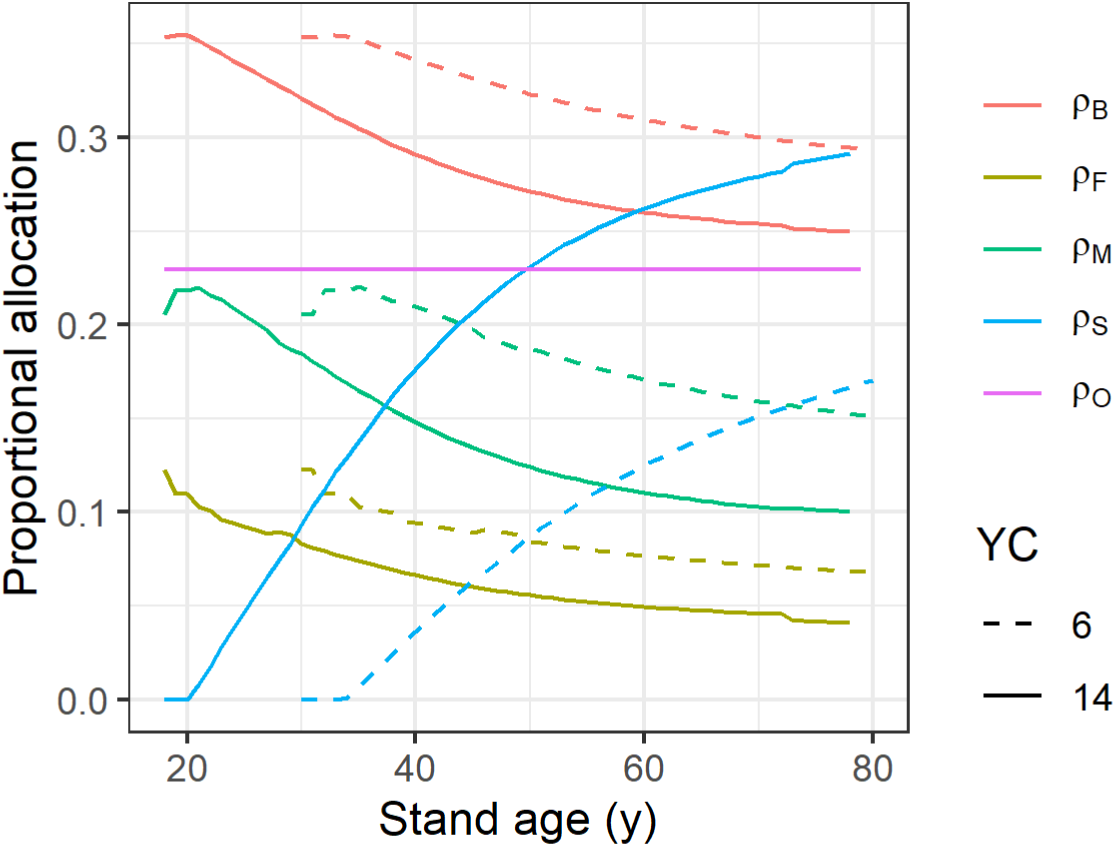
The proportional allocation of above ground biomass to biofuels (*ρ*_*B*_), fast (i.e. paper, *ρ*_*F*_), medium (i.e. panelling, *ρ*_*M*_) and slow (i.e. sawn wood, *ρ*_*S*_) decaying wood products, and in situ off cuts (*ρ*_*O*_) predicted by CARBINE for Sitka spruce stands of yield class 6 and 14. The minimum age available for a given yield class reflects the minimum age at which yield classes can be empirically distinguished.

The unitless decay efficiency, *δ*_*X*_, represents the proportion of the biomass within a given product or tissue type that ultimately ends up as an atmospheric carbon. A proportion (1 − *δ*_*X*_) remains fixed in an implicit, non-decaying pool e.g. soil organic carbon in anaerobic zones of the peat. Parameterising *δ*_*X*_is non-trivial, not least because the size of any ‘fixed’ carbon pool will depend on chemical and hydrological factors as well as the temporal extent of the analysis. Therefore, we set *δ*_*X*_ = 1 for all *X* by default, i.e. assuming that 100% of the forest biomass ultimately decays to atmospheric CO_2_, but leave the option for the user to relax this assumption if required.

Carbon stocks are represented using a combination of 3PG and CARBINE since 3PG is a more sensitive model but, as implemented here (Eqs. (3) – (9)), lacks the necessary partitioning of carbon between tissue types. The total carbon stored in forest biomass at harvesting, *C*_*tot*_(*t*_ℎ*arv*_) (t C ha^-1^), is predicted from 3PG by the cumulative sum of net primary productivity from time *t* = 0 to the harvesting age *t*_ℎ*arv*_:

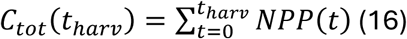

The carbon stored in roots, stem, branches and foliage, *C*_*r*_(*t*_ℎ*arv*_), *C*_*s*_(*t*_ℎ*arv*_), *C*_*b*_(*t*_ℎ*arv*_) and *C*_*f*_(*t*_ℎ*arv*_) (t C ha^-1^) respectively, is assessed by rescaling *C*_*tot*_(*t*_ℎ*arv*_) by unitless ratios output by CARBINE.

If forest material is left to decompose on site, initial decomposable carbon stocks *S*_*X*ℎ*arv*_are given directly from *C*_*X*_(*t*_ℎ*arv*_) e.g. initial decomposable stem biomass (t CO_2_ eq.) is given by *S*_*S*ℎ*arv*_ = *C*_*CO*2−*C*_*δ*_*S*_*C*_*s*_(*t*_ℎ*arv*_) etc. This initial carbon stock then decays to the atmosphere according to Eq. (12) – (13). If the user chooses to model mulching of above-ground forestry biomass, the stem and branch biomasses are combined in a single compartment with a half-life corresponding to mulched material. Note that even when the forestry is harvested and processed into wood products, some biomass is left to decay on the site: all the root and foliage biomass and a proportion of the woody biomass representing in situ processing losses (off-cuts) which are considered as mulch. The half-lives for the various decay compartments are given in Tab. 3.

**Tab. 3.** Half-lives of the various decay products.

| Wood product/tissue type | Half-life (yr) | Source |
| --- | --- | --- |
| Paper | 2 | Domke et al. (2019) |
| Panel | 25 | Domke et al. (2019) |
| Sawn timber | 35 | Domke et al. (2019) |
| Foliage | 1.23 | CARBINE technical guide (Forest Research, 2025) |
| Roots | 19 (16 – 23) | Olajuyigbe et al. (2011) |
| Branches | 2.79 | CARBINE technical guide (Forest Research, 2025) |
| Stem wood (logs and stumps) | 13 (12 – 14) | Olajuyigbe et al. (2011) |
| Mulched wood | 2.79 | CARBINE technical guide (Forest Research, 2025) |

#### 2.6.3. Emissions from biofuels

The PEATREST LCA accounts for emissions associated with the conversion of forest biomass to biofuels. The proportion of forest biomass allocated to biofuels, *ρ*_*B*_(*t*_ℎ*arv*_), is an additional output from CARBINE (Fig. 4). When biomass is burnt, all of the carbon stored in this material, *S*_*B*ℎ*arv*_ = *C*_*CO*2−*C*_*ρ*_*B*_(*t*_ℎ*arv*_)*C*_*a*_(*t*_ℎ*arv*_), is immediately lost to the atmosphere. For simplicity, all biofuels are assumed to be consumed in the harvesting year such that biofuel emissions represent a single one-off contribution to the carbon accounting. The energy generated (MWh) by burning biofuels offsets energy from the national grid. Thus, the carbon losses associated with burning forest biomass are offset by emissions avoided from alternative fuel sources.

The net emissions associated with the use of biofuels, *L*_*B*_ (t CO_2_ eq.) are given by

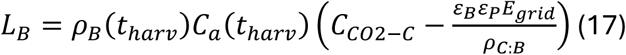

As above the Carbon: Biomass ratio *ρ*_*C*:*B*_ converts the estimated carbon stock into a biomass (t Biomass ha^-1^). *ε*_*B*_ is the power-value of the forestry biomass set to 3.61 MWh t-1, (Forest Research^1^), which makes the simplifying assumption that the moisture content of the biofuel is 30%. *ε*_*P*_is the unitless conversion efficiency of the biofuel power plant set to 0.352 (indicative value from the Port Talbot CFBC biomass plant, McIlveen-Wright et al. 2013). *E*_*grid*_is the grid-mix emissions factor (0.225 t CO_2_ MWh^-1^, GOV.UK^2^).

The emissions factor for biofuels (CO_2_ emissions divided by energy generated) based on these default parameters is *C*_*CO*2−*C*_*ρ*_*C*:*B*_*ε*_*B*_^−1^*ε*_*P*_^−1^ = 1.44 t CO_2_ MWh^-1,^ i.e. around 6 times greater than for the grid mix. Note that combustion emissions associated with biofuels are frequently excluded from analyses of this type under the assumption that at long enough timescales all non-fossilised biomass ultimately returns to gaseous form such that burning produces no net change in biogeochemical concentrations (e.g. Bates and Henry, 2009). Here we *do* consider emissions from biomass combustion since the temporal scope of the LCA is likely to be of the order of decades, i.e. much less than the time required for forests to release their carbon via natural mechanisms.

#### 2.6.4. Emissions from transportation of wood products

Transport emissions are accounted for when forest biomass is processed into wood products including biofuels. Emissions are estimated by multiplying the biomasses transported, *B*_*X*_(*t*_ℎ*arv*_) = *ρ*_*C*:*B*_^−1^*ρ*_*X*_(*t*_ℎ*arv*_)*C*_*s*_(*t*_ℎ*arv*_) (t Biomass), by the user input distances to processing sites and the emissions factor *E*_*transport*_ = 39.33 (38.5 – 40.15) g CO2 km^-1^ t^-1^ assuming fully loaded trucks of max load around 20 t (Morison et al. 2012).

### 2.7. Restoration of peatland ecosystem function

The restoration of peatland ecosystem function is represented by first computing the emissions rates a) following removal of the trees and b) once ‘full’ ecosystem function has returned. Water tables depths have been shown decrease by around 45% following removal of trees from afforested peatlands (Gaffney et al. 2018), likely because of the cessation of evapotranspiration and the interception of rainwater by the canopy. Thus, the LCA rescales the user input water table depth prior to harvesting by a corresponding amount to represent the post-harvesting, pre-restoration hydrology. Pristine blanket bogs typically have a median annual water table depth of around 4 (1-6) cm (Gaffney et al. 2018, Hancock et al. 2018), a depth at which peats are predicted to be net carbon sinks (Evans et al. 2021). Post-restoration water tables are set to 5 (1-10) cm accordingly. Pre-and post-restoration carbon fluxes for these water table depths are modelled using Eqs. (1) and (2).

To avoid discontinuities in the emissions dynamics, and, crucially, to explore the intermediate dynamics in carbon mitigation effects, pre- and post-restoration emissions rates are mapped to an arbitrary asymptotic function parameterised by two key inputs: the restoration time and the shape of the restoration trajectory.

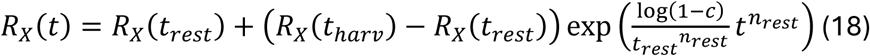

Here *R*_*X*_(*t*_ℎ*arv*_) and *R*_*X*_(*t*_*rest*_) are the pre- and post-restoration emissions rates respectively, *t*_*rest*_is the time to restoration of ‘full’ ecosystem function and *n*_*rest*_is a shape parameter controlling the lag in the restoration trajectory (Fig. 5). *c* is a convergence limit set to 0.999, required since the function formally converges at *t* = Inf.

**Fig. 5.**
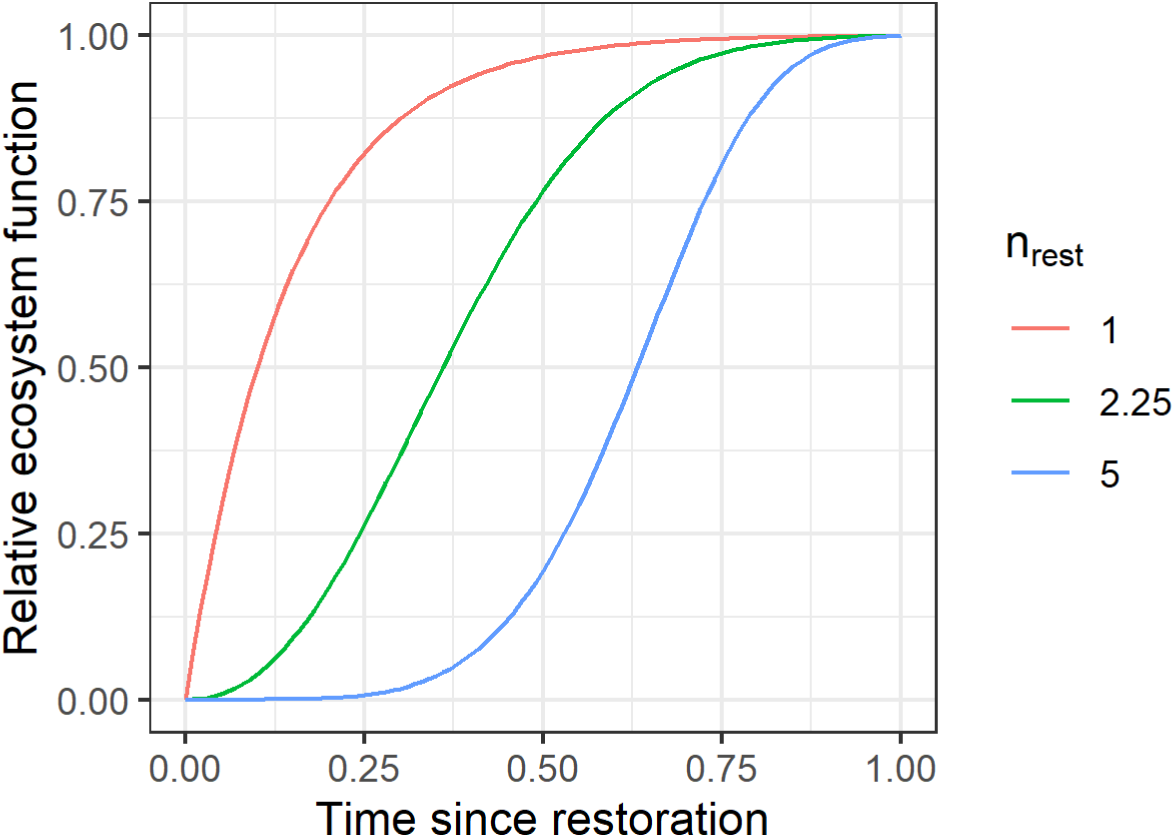
The arbitrary asymptotic function (Eq. (18)) used to represent the restoration of ecosystem function through time. Here the pre-and post-restoration emissions rates are shown in relative units (i.e. set to 0 and 1 respectively), while the restoration time is set to 1. The lag in the restoration trajectory in controlled by the shape parameter *n*_*rest*_.

### 2.8. Carbon payback time

Two different metrics of carbon restoration time are computed from the composite time series: the flux intercept *t*_*flux*_, and the carbon payback time *t*_*payback*_. In practice *t*_*flux*_is the time following harvesting at which the peatland sequestration time series first intercepts the forestry sequestration time series. *t*_*payback*_is the intercept of the *cumulative* sequestration trajectories.

### 2.9. Scenario modelling

To demonstrate model outputs below, two sites with contrasting management strategies are represented. Note that model inputs are arbitrary and selected only for illustrative purposes. When implemented to support restoration practitioners, inputs will be set based on a combination of expert knowledge, monitoring and modelling.

Area 1 is harvested and forest biomass removed for processing into wood products. In line with empirical results (Rydgren et al., 2025), restoration time is set to *t*_*rest*_ = 50 years with *n*_*rest*_ = 1.5 roughly reproducing an observed trajectory in water table depth recovery (Gaffney et al., 2018). For Area 2, forestry is felled to waste with biomass left to decay on site. We make the crude assumption that un-mulched timber left on site doubles restoration times to *t*_*rest*_ = 100 years and causes a lag in the restoration trajectory corresponding to a value of *n*_*rest*_ = 4 since slow-decaying stem biomass will offer little nutrient input and delay the growth of bog plants by passively competing for space and light. For both areas, restoration interventions include drainage damming, bund creation and peat smoothing/stump flipping.

For each area the model is run three times, with yield class and post-restoration water tables modified to give minimum, expectation and maximum values for the carbon mitigation times. Yield classes were set to 6, 10 and 14, while restored water tables were 0.01, 0.05 and 0.1 m. All other model inputs were fixed. A summary of the key model inputs corresponding to Areas 1 and 2 is given in Appendix D.

### 2.10. Sensitivity analysis

Sensitivity analysis is used to explore which parameters need to be estimated with the greatest accuracy, though note that it can also highlight which management strategies are most likely to substantially impact mitigation times. The sensitivity of the model outcomes is assessed using an iterative simulated experiment taking the inputs for Area 1 (with yield class and post-restoration water tables set to 8 and 0.05 m respectively) as the initial parameterisation.

Each model input in turn was modified by multiplying its expected value by a scaling variable in the range 0.2 to 2.0. All other inputs were kept fixed. The model was then evaluated and *t*_*flux*_and *t*_*payback*_estimated. Note that the sensitivity analysis does *not* explore the full parameter space of the model. Instead, it represents variation in model outcomes in the vicinity of a single vector within the parameter space corresponding to the expected values for Area 1. A full scan of the parameter space of the model would be almost impossible to interpret given the number of parameters that would need to be considered however, repeating the analysis with the yield class set to 6 or 10 produced similar results.

The sensitivity of model outcomes to a given input was assessed as the slope of the linear regression of the relative change in *t*_*flux*_or *t*_*payback*_as a function of the relative change in the input. The model frequently responds non-linearly to variation in inputs. As such the linear slope gives a crude but informative representation of the sensitivity.

## 3. Results and Discussion

In the following figures, sequestration and emissions are typically given in units t CO_2_ eq. Absolute quantities (t CO_2_ eq.), are used instead of rates (t CO2 eq. ha^-1^ yr^-1^) since *t*_*payback*_ is defined by reference to total carbon storage (t CO_2_ eq.). Here the values are equivalent since both areas have a spatial extent of 1 ha, and the timestep size is 1 year.

### Counterfactual (forestry left in situ)

The compound time series (Fig. 6, black) generated by summing sequestration by the forestry (Fig. 6, red), emissions from the peat under the trees (Fig. 6, green) and leaching of aquatic carbon (Fig. 6, blue) is frequently negative, implying woodland that is a net carbon source. In the case of low yield class forestry, emissions from the soils exceed sequestration even at peak productivity, such that the woodland is *never* a net carbon sink. Encouragingly, the model appears to conform to the rule-of-thumb expectation that forestry planted on drained peatland becomes a net carbon sink between YC 6 and YC 10 (YC 8, Anderson 2020).

**Fig. 6.**
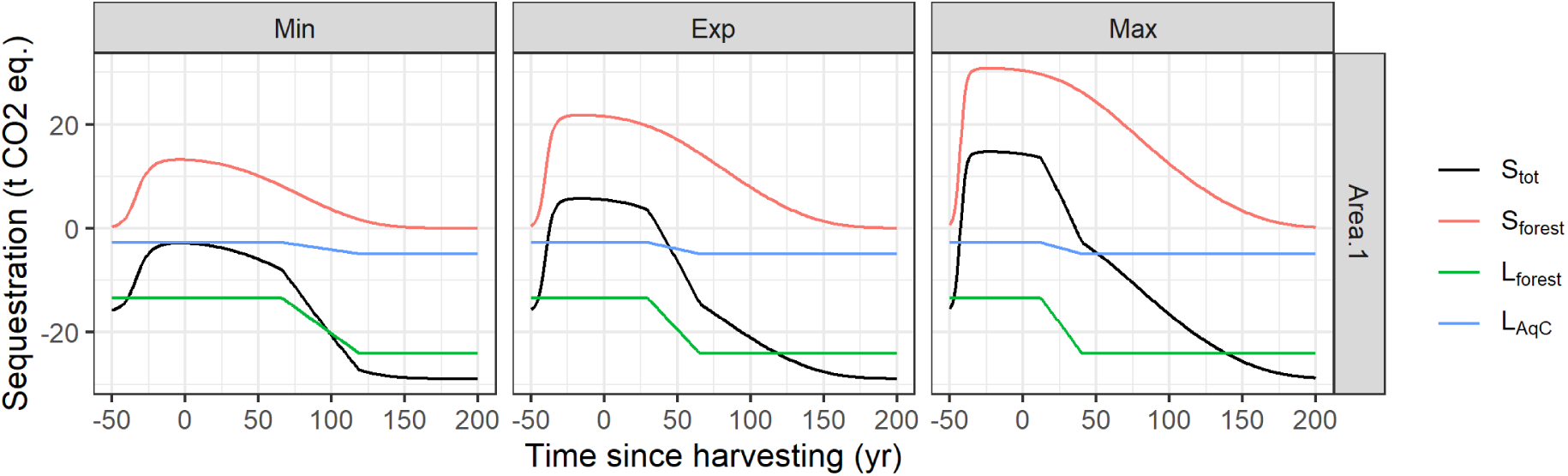
Net sequestration by the forestry (black) is given by the balance between forestry NPP (red), CO_2_ + CH_4_ emissions from the drained peats under the trees (green) and losses of aquatic carbon due to leaching (blue). Low yield class forestry (Min: YC 6) may be a net carbon source even at peak forest sequestration. Areas 1 and 2 are identical with respect to the counterfactual.

In the model evaluation show, drainage and maximum rooting depth were kept fixed such that minimum and maximum carbon loss rates from soils are identical for all yield classes. Yield class does, however, impact the rate of accumulation of root biomass. Therefore, high yield class forestry reaches maximum root depth earlier. This effectively reduces the duration of peak net sequestration for high yield class forestry, a phenomenon that could be further explored in empirical studies. This narrowing of the peak reduces the total carbon storage by the forestry and therefore impacts carbon mitigation times.

### Continuous-time emissions due to forest biomass decay

Carbon emissions due to the decomposition of forest biomass increase with yield class (greater total biomass) but are also sensitive to management strategy (Fig. 7). The half-lives of slow and medium lived wood products are lower than those of stem wood (Tab. 3). As such more carbon is emitted in the years immediately after harvesting when forestry biomass is removed from site and processed. For this model run, 100% of the forest biomass ultimately ends up as atmospheric carbon (*δ*_*X*_ =1 for all *X*) meaning differences in decay emissions are strictly transitory. In reality, some biomass will enter very slow decaying pools (e.g. in anaerobic zones of rewetted peats). Estimating the size of such fixed carbon pools may improve the sensitivity of the LCA to different management scenarios.

**Fig. 7.**
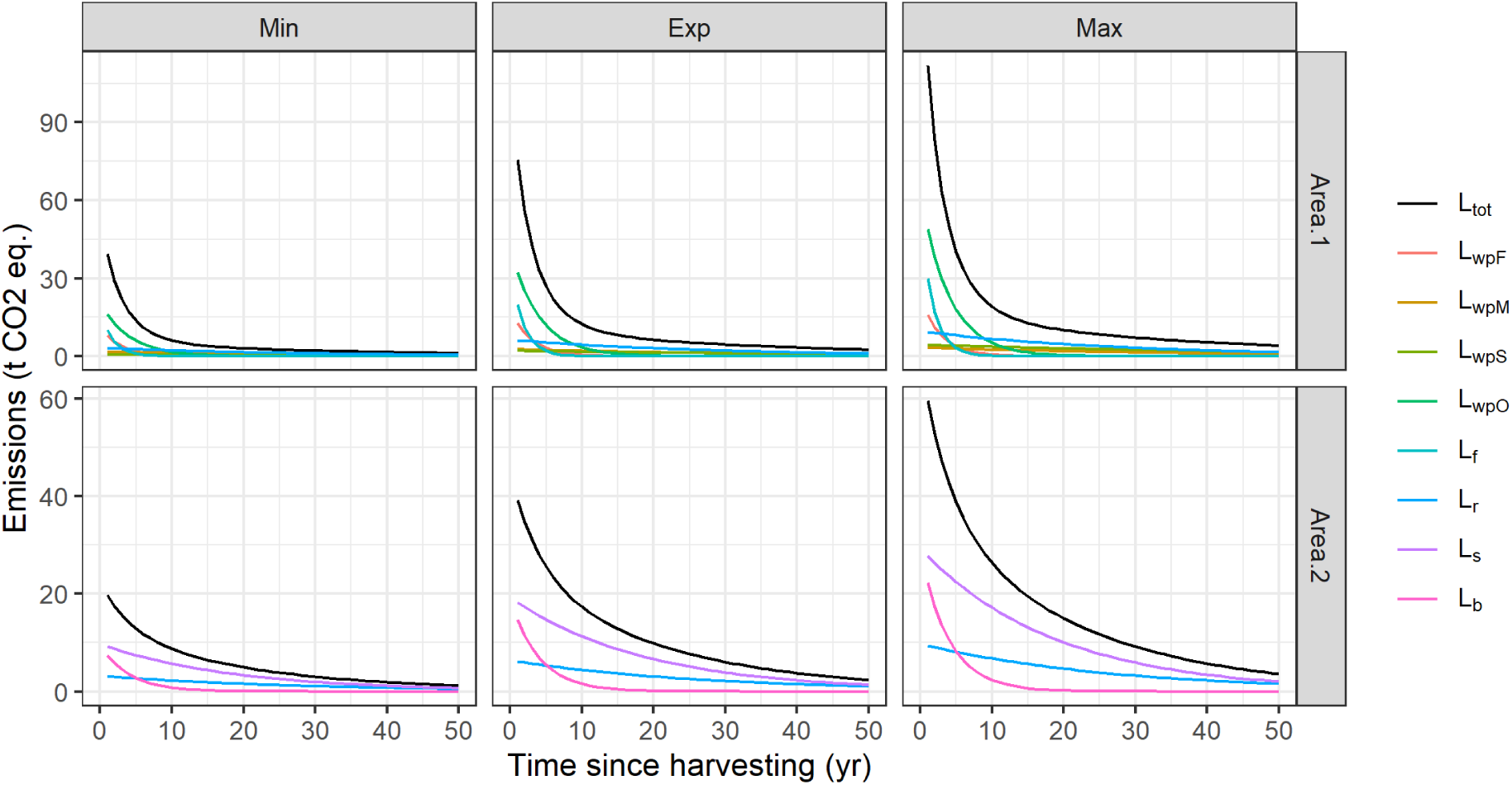
Emissions due to the decay of forest biomass, either converted into wood products (Area 1), or left on site (Area 2). Total emissions (black) are substantially greater at intermediate timescales when biomass is processed into wood products due to the relatively short half-lives of the processed material. *wpX* denotes wood product *X* (Fast, Medium, Slow decaying or in situ processing losses ie. Offcuts). Subscripts *f*, *r*, *s*, and *b* refer to foliage, root, stem and branch biomass pools respectively.

### Discrete time emissions due to management interventions

One-off emissions due to management and restoration (Fig. 8) are dominated by the burning of biofuels and harvesting (or mulching, if selected by the user), though peat smoothing/stump flipping also represents a substantial contribution. Emissions due to transport and hydrological restoration (damming and bunding) are much smaller. Note, energy intensive restoration practices like re-turfing (active intervention establishing bog vegetation) were not included in this evaluation of the model.

**Fig. 8.**
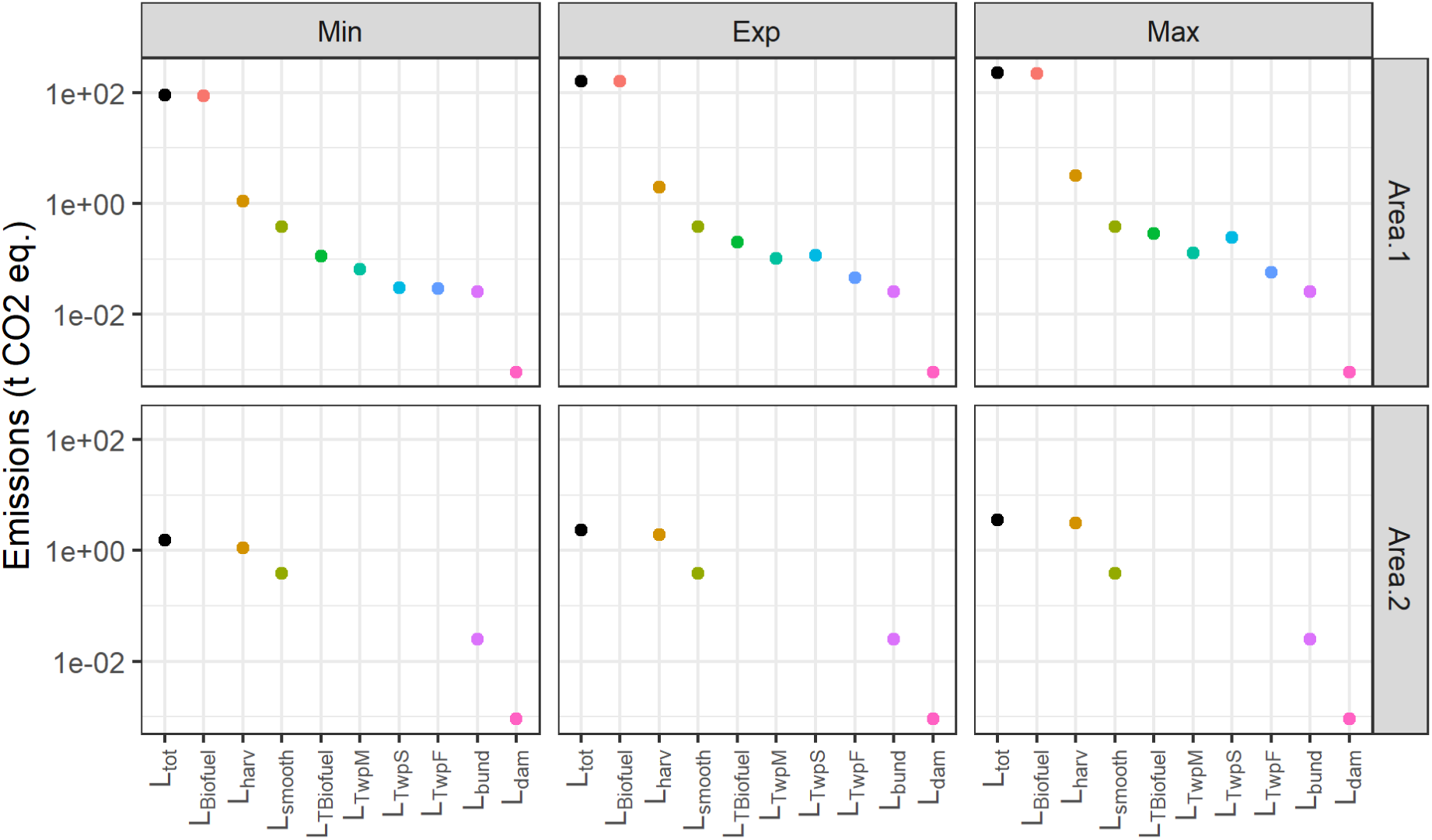
Discrete emissions due to harvesting and restoration activities and the burning of biofuels. Biofuel burning and harvesting emissions dominate the management emissions (note the logarithmic y axis). The subscript *T* on the x-axis indicates transport emissions.

### Peatland carbon sequestration dynamics

The restoration of peatland ecosystem function is shown in Fig. 9. Area 1 becomes a net carbon sink after 17-27 years (dashed lines, Fig. 9). Area 2 becomes a net carbon sink 65-79 years after harvesting. In accordance with Eqs. (1) and (2), the maximum sequestration rate depends sensitively on the post-restoration water table depth. Note the time to full restoration of ecosystem function was set to 50 and 100 years respectively. The fact that restoration of net zero GHG emissions coincides approximately with empirical observations (Hambley et al. 2018, Fundira et al. 2025) reflects the choice of inputs rather than an emergent prediction from the model dynamics. However, it does highlight the benefits of including a non-linear trajectory instead of a simple step function to represent the restoration of ecosystem functionality.

**Fig. 9.**
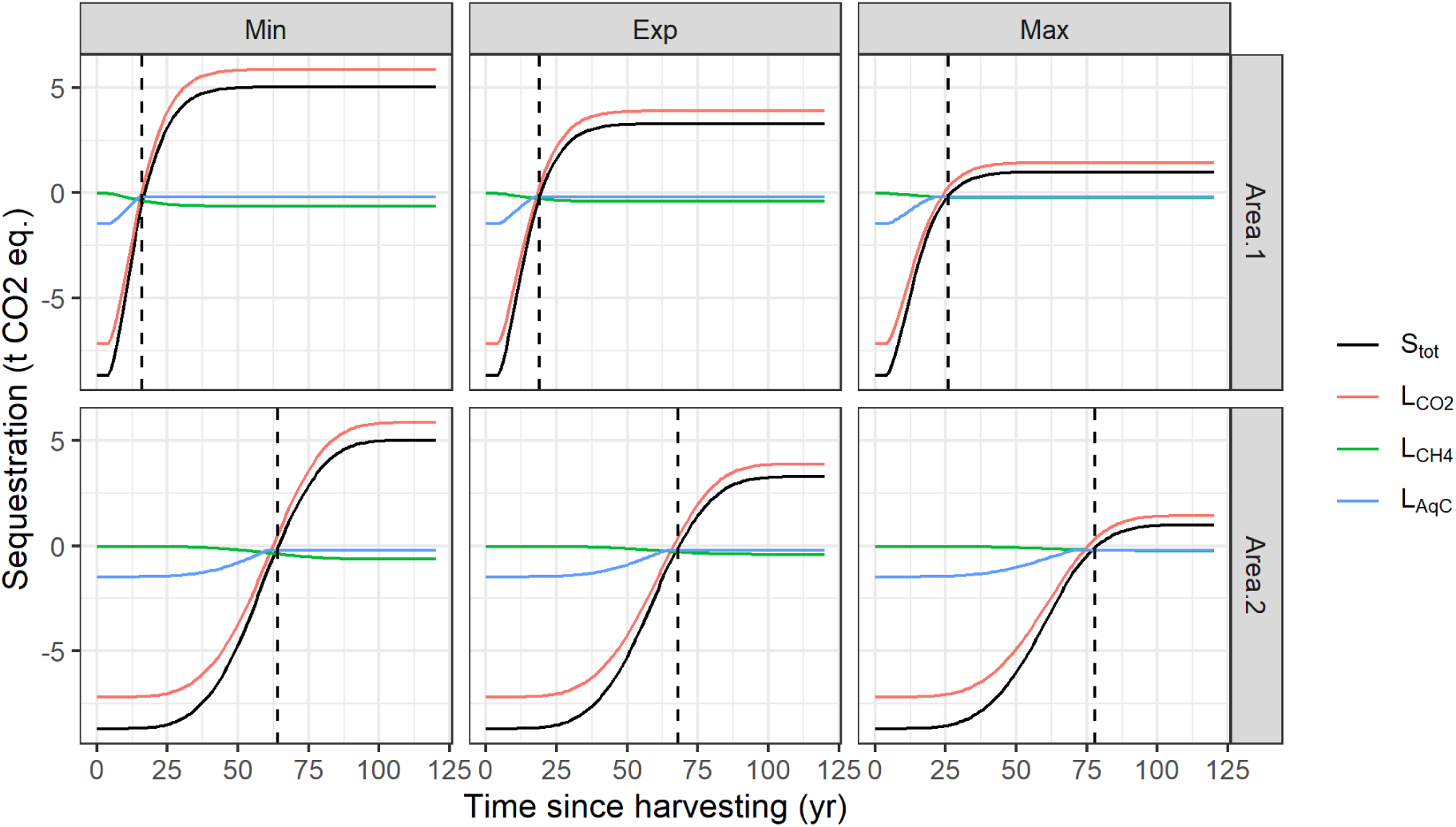
The restoration of peatland sequestration (black) represented by pre- and post- restoration emissions rates mapped to an arbitrary asymptotic function (red and green). Aquatic carbon losses (blue) are generated by rescaling total gaseous emissions down to a minimum value corresponding to observed values for undrained peatlands. The restoration trajectory is defined by user input restoration times and post-restoration water table depths, set here to 0.01, 0.05, and 0.10 m for Min, Exp, and Max respectively.

### Flux intercept and Carbon payback time

The peatlands are first predicted to sequester carbon at a faster rate than the forestry would have *t*_*flux*_ = 18-59 years after harvesting (see Tab. 4 and Fig. 10) depending on management strategy, yield class and post restoration water table. *t*_*flux*_ was higher for Area 2 due to the assumed greater restoration time when forest biomass is left un-mulched on the surface of the peat. Modelling results suggest that peatland restoration provides a net *mitigation* benefit within a few decades of deforestation and restoration when plausible inputs are used. Note that *t*_*flux*_can occur when the peatland is still a net carbon source due to the tendency for forestry emissions to exceed sequestration (Fig. 10). This again highlights the importance of including a restoration trajectory in the model dynamics. If the restoration of peatland ecosystem function is represented using a simple step function (Evans et al. 2023), mitigation times are strictly greater or equal to the user input restoration time.

**Fig. 10.**
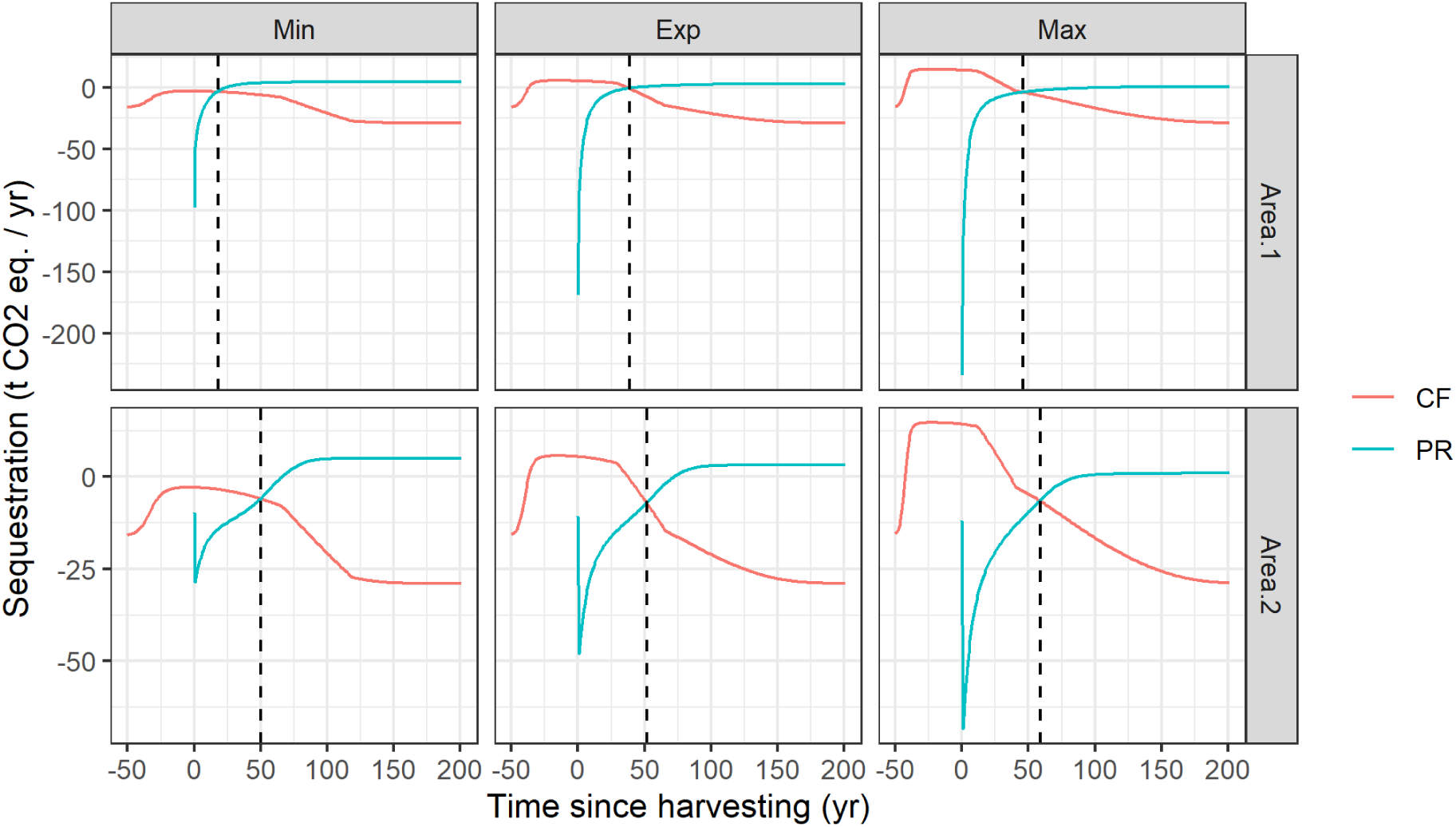
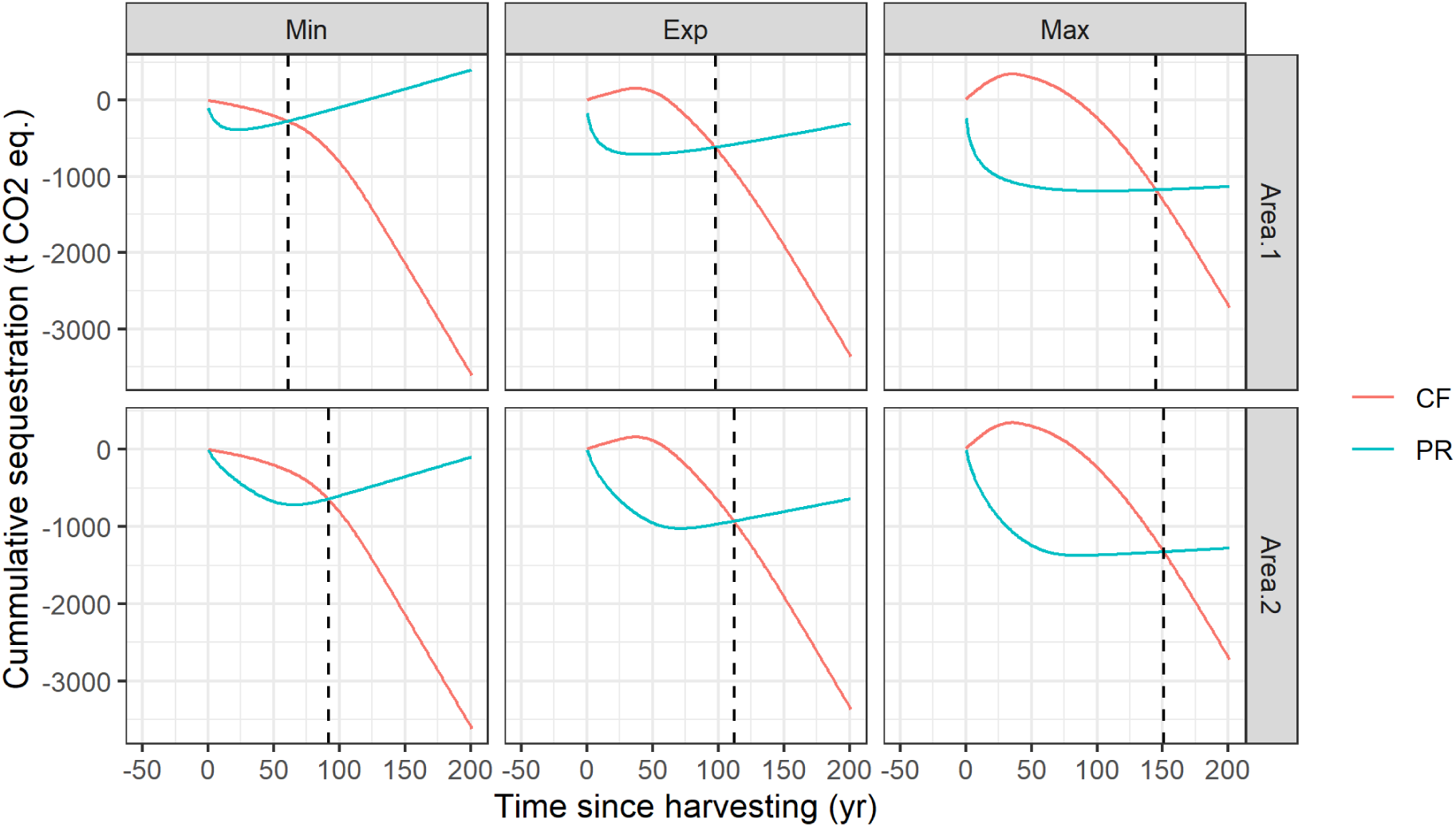
The carbon flux intercept (above, dashed vertical line) and the carbon payback time (below, dashed vertical line) assessed via the intercept of the compound time series of sequestration rates (above) or carbon storage (below). The counterfactual (CF, red) and the peatland restoration scenario (PR, blue) are shown.

**Tab. 4.** The carbon flux intercept *t*_*flux*_ and payback times *t*_*payback*_ for the two management strategies (areas).

|  | Area 1 |  | Area 2 |  |
| --- | --- | --- | --- | --- |
| | $t_{flux}$ (yr) | $t_{payback}$ (yr) | $t_{flux}$ (yr) | $t_{payback}$ (yr) |
| Min (YC 6, $W_{rest}$ 0.01) | 18 | 61 | 50 | 92 |
| Exp (YC 10, $W_{rest}$ 0.05) | 39 | 98 | 52 | 112 |
| Max (YC 14, $W_{rest}$ 0.10) | 46 | 145 | 59 | 151 |

Carbon payback times varied from *t*_*payback*_ = 61-151 years after harvesting, with Area 2 again the slower of the two. *t*_*payback*_tends to occur during the phase in which the peatland has yet to pay back the carbon debt incurred during the early phase of the restoration. The model suggests that the time following harvesting before peatland carbon storage becomes *net positive* could be multiple centuries. This highlights two important points. First is that the long-term implications of peatland restoration in terms of carbon storage depend sensitively on the speed of hydrological restoration. If deforested peatlands are left dry for many years, the accumulated carbon debt could be very large. The second is that the net gains associated with restoration also depend sensitively on the end use of the forestry biomass since much of the carbon debt can be attributed to emissions from fast decaying wood products and biofuels. Carbon payback times could be substantially reduced via careful management of harvested forestry biomass.

### Sensitivity analysis

The outcome of the sensitivity analysis is shown in Fig. 11. Only inputs that produce a change of at least 1 year in the mitigation times are shown. The actual variation in model outcomes with variation in inputs is shown in Appendix E (Figs. E1 and E2). These highlight the tendency for non-linearities in the sensitivity to model inputs. The carbon flux intercept *t*_*flux*_is sensitive to fewer inputs than the carbon payback time *t*_*payback*_. This reflects the fact that the sequestration trajectory retains no ‘memory’ of the early phase of the restoration process, while net carbon storage, the cumulative sum of the sequestration trajectory, does.

**Fig. 11.**
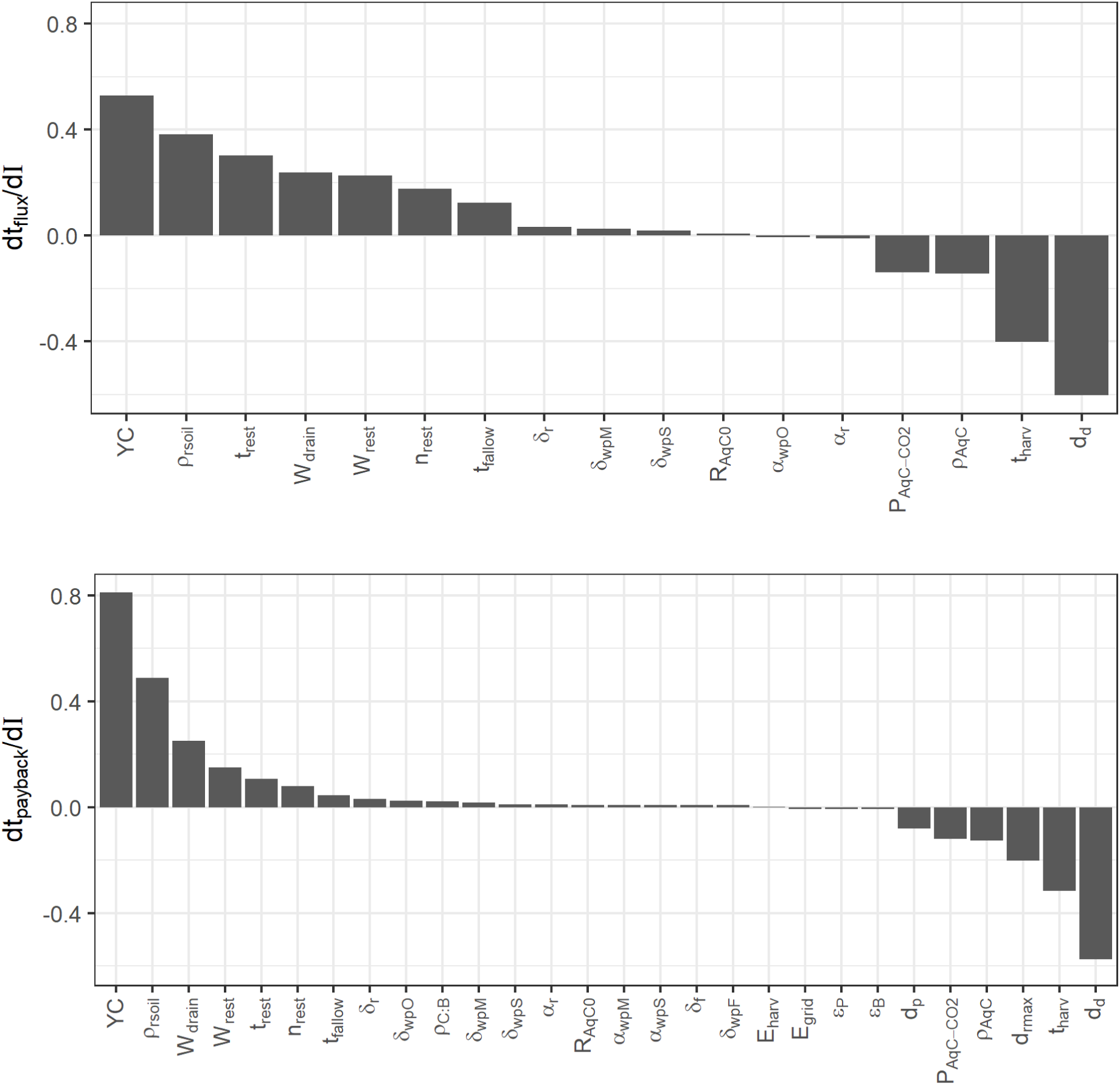
The linear slopes of the proportional change in carbon flux intercept (above) and carbon payback time (below) for a proportional change in each input variable *I*. Note that a value of 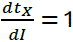 implies that doubling the input variable *I* leads to a doubling in the mitigation time.

The most sensitive inputs are broadly consistent for the two metrics. Yield class (YC), root zone biomass density (*ρ*_*r*,*soil*_), pre-restoration water table depth (*W*_*drain*_), post-restoration water table depth (*W*_*rest*_), restoration time and shape parameter (*t*_*rest*_ and *n*_*rest*_) and the length of the fallow period between harvesting and restoration (*t*_*fallow*_) all correlate positively with mitigation times. Negative associations are found with the harvesting age (*t*_ℎ*arv*_), the parameters of the aquatic carbon loss sub-model (*ρ*_*AqC*_ and *P*_*AqC*−*CO*2_), peat, rooting and drainage depths (*d*_*p*_, *d*_*r*.*max*_ and *d*_*d*_).

The sensitivity to YC implies that accurate parameterisation of the 3PG module is a crucial component of the model. Care should be taken to ensure that the productivity of the forestry predicted by 3PG matches empirical expectation and where the counterfactual deviates from reasonable expectation, model inputs should be modified accordingly.

The biomass density of the rooting zone *ρ*_*r*,*oil*_is a sensitive parameter since it controls the rate at which max root depths are attained and therefore impacts the emissions from drained peats under the forestry. Notably, however, it only strongly impacts model outcomes when set to a value of much less than the default (Figs. E1 and E2). In the empirically observed range (0.039 – 0.08 t Biomass m^-3^, Nicoll et al. 2006), it’s impact on mitigation times is limited suggesting the high sensitivity to *ρ*_*r*,*soil*_ is not a major concern.

Pre-restoration water table depths *W*_*drain*_are estimated by rescaling the user input water table under the trees by an empirically observed factor (45% decrease following tree removal). This proportion should be updated as additional results become available. Post-restoration water table depths *W*_*rest*_unsurprisingly impact predicted mitigation times. Note that the non-linearity in the response of *t*_*flux*_and *t*_*payback*_to *W*_*rest*_(Figs. E1 and E2) suggest that for *W*_*rest*_ ≤ 0.1 m, results are broadly similar partly due to the onset of methanogenesis. While minimising the water table depth following restoration should always be a priority, this suggests the range 0 < *W*_*rest*_ ≤ 0.1 m will produce broadly similar net carbon benefits (around a 10% difference in mitigation times).

Restoration timescales *t*_*rest*_and *n*_*rest*_are clearly vital inputs. Developing monitoring and modelling programs to constrain these estimates is a clear research priority. Surprisingly, there is not a 1:1 association between either *t*_*rest*_or *t*_*fallow*_and the mitigation times (Fig. 11, slopes < 1). This is because for each additional year the peatland remains drained, the counterfactual predicts additional losses from the forestry.

Varying harvesting age *t*_ℎ*arv*_has a complex impact on model outcomes, in particular for *t*_ℎ*arv*_ < 50 (Figs. E1 and E2). The removal of younger stands typically means an increase in the lost sequestration potential as well as a lower hydrological impact on the soils under the trees. Meanwhile, greater total yield in more mature stands means more carbon that ultimately returns to the atmosphere as forest biomass decays, increasing payback times. Additional complexities arise due to the decrease in the proportion of forest biomass allocated to biofuels for older stands. As such, it is not possible to generalise the sensitivity of the model to *t*_ℎ*arv*_but further exploration of the model could provide rule of thumb estimates for minimum harvesting ages in order to achieve carbon mitigation in relevant timescales.

Model outcomes are sensitive to the parameters defining the rate of aquatic losses to atmosphere *P*_*AqC*−*CO*2_and *ρ*_*AqC*_. These should also therefore be updated as estimates are improved. In future a robust, process- or regression-based model for aquatic carbon losses would be desirable.

Increasing the drainage depth *d*_*d*_increases emissions from the soils under trees, especially in young stands. This reduces payback times by reducing the carbon storge capacity of the forestry. Increasing peat depth *d*_*p*_up to the maximum rooting depth decreases payback times since the effective water table depth is set by the minimum of these two values. In practice, however, peat depths less than root depths are unlikely, in particular if the lower soil strata are not amenable to root growth, therefore this sensitivity is potentially a model artifact.

The decay rate parameters, including *δ*_*X*_, impact outcomes but are typically not very sensitive meaning coarse estimates or crude assumptions are likely to be sufficient for these inputs. Payback times strictly increase with *δ*_*X*_ since *δ*_*X*_ linearly scales the total carbon available for decomposition to the atmosphere. The correlation between *ɑ*_*X*_and mitigation times can be either positive or negative. The sign of the relationship depends on the size of the carbon pool and the expectation value of the input.

## 4. LCA caveats and conclusions

The sensitivity analysis highlights which inputs/sub-models are most important to determining model outcomes. Clearly, the model must be caveated by noting that predictions will be accurate only insofar as the sensitive parameters are well constrained. As additional studies of peatland biogeochemical processes are undertaken, these model parameters should be updated accordingly.

The emissions rates sub-model, Eqs. (1) and (2), was parameterised by considering average annual water table depths (Evans et al. 2021). In practice, intra-annual variation in hydrology may impact emissions rates. This effect will be particularly relevant in the case that water table depths are a) asymmetrically distributed around annual averages and b) emissions rates respond non-linearly to hydrological conditions. Additionally, the counterfactual projects emissions from drained peats forward multiple decades. These projections ignore the fact that carbon stocks in drained peats will eventually be depleted. Ideally, a decaying function should be used to represent emissions from peats for which carbon stocks are not being replenished.

The time required for restoration of ecosystem function is a crucial unknown that this model cannot address. It is vital, therefore, that it be understood as a tool developed in parallel to monitoring and modelling studies that aim to shine a light on the typical timescales for ecological restoration. The LCA does, however, consider a suite of processes both local to and distant from the restoration project when assessing the carbon mitigation times. In this, it represents an important step change in the quantitative modelling of carbon fluxes along peatland restoration gradients.

Finally, while rigorous quantitative validation of the model is impossible without long-term experimental data against which to compare predictions, the LCA appears to capture the rule-of-thumb that forestry planted on drained peats becomes a net carbon sink at around yield class 8 (Anderson, 2020). Thus, despite the caveats noted above, this provides an early quantitative test of the predictions generated using the PEATREST LCA.

## Data availability statement

All equations required to reproduce the analysis are included in the main text of the article. The R code used to run the model will be publicly available in a GitHub repository on publication. All model parameters are either explicitly stated in the manuscript or included in input files in the GitHub repository.

## CRediT authorship contribution statement

**Jacob O’Sullivan:** Conceptualization, Formal analysis, Investigation, Methodology, Software, Validation, Visualization, Writing – original draft. **Carly Whittaker:** Data curation, Writing – original draft. **George Xenakis:** Funding acquisition, Writing – review & editing. **Toby Robson:** Writing – review & editing. **Mike Perks:** Conceptualization, Funding acquisition, Supervision, Writing – review & editing.

## Declaration of interests

The authors declare the following financial interests/personal relationships which may be considered as potential competing interests: All authors report financial support was provided by United Kingdom Department for Environment Food and Rural Affairs.

1 https://www.forestresearch.gov.uk/tools-and-resources/fthr/biomass-energy-resources/reference-biomass/facts-figures/potential-yields-of-biofuels-per-ha-p-a/

2 https://www.gov.uk/government/publications/greenhouse-gas-reporting-conversion-factors-2024

3 Forestry mulchers: 4 solutions on the market right now | Forestry Journal

4 Forestry mulchers with fixed hammers for tractor | Rabaud

5 Mulching - RGL Forestry

6 PT-300.pdf

7 FTX300 Mulching Tractor / Forestry Mulcher / Tracked Mulcher

## Appendix

### A. Full list of variables used in the PEATREST LCA

**Tab. A1.** Full list of variables used in the PEATREST LCA.

| Variable | Symbol | Units | Variable | Symbol | Units |
| --- | --- | --- | --- | --- | --- |
| **Emissions rates modelling** | | | Ratio of total aquatic carbon to total gaseous carbon loss | $\rho_{AqC}$ | - |
| CO<sub>2</sub> emissions rate | $R_{CO_2}(t)$ | t CO<sub>2</sub> ha<sup>-1</sup> yr<sup>-1</sup> | Empirical aquatic carbon loss from peatlands | $R_{AqC0}$ | t CO<sub>2</sub> eq. ha<sup>-1</sup> yr<sup>-1</sup> |
| CH<sub>4</sub> emissions rate | $R_{CH_4}(t)$ | t CO<sub>2</sub> eq. ha<sup>-1</sup> yr<sup>-1</sup> | Proportion of aquatic carbon that ends up as atmospheric CO<sub>2</sub> | $P_{AqC-CO_2}$ | - |
| Units C to units CO<sub>2</sub> conversion | $C_{CO_2-C}$ | units CO<sub>2</sub> (units C)<sup>-1</sup> | **Harvesting / management modelling** | | |
| Water table depth | $W(t)$ | m | Harvested area | $A_{harv}$ | ha |
| CO<sub>2</sub> emissions | $L_{CO_2}(t)$ | t CO<sub>2</sub> | Stand age at harvesting | $t_{harv}$ | yr |
| CH<sub>4</sub> emissions | $L_{CH_4}(t)$ | t CO<sub>2</sub> eq. | Transportation emissions factor | $E_{transport}$ | g CO<sub>2</sub> km<sup>-1</sup> t<sup>-1</sup> |
| Total gaseous emissions | $L_{Tot}(t)$ | t CO<sub>2</sub> eq. | Harvesting emissions factor | $E_{harv}$ | kg CO<sub>2</sub> m<sup>-3</sup> |
| **Forest growth modelling (3PG)** | | | Mulching emissions factor | $E_{mulch}$ | kg CO<sub>2</sub> ha<sup>-1</sup> |
| Absorbed photosynthetically available radiation | $\phi_{pa}(t)$ | MJ m<sup>-2</sup> yr<sup>-1</sup> | Bund creation emissions factor | $E_{bund}$ | kg CO<sub>2</sub> ha<sup>-1</sup> |
| Annual average radiation above the canopy | $\phi_p$ | MJ m<sup>-2</sup> yr<sup>-1</sup> | Drainage damming emissions factor | $E_{dam}$ | kg CO<sub>2</sub> ha<sup>-1</sup> |
| Light extinction coefficient | $k_{ext}$ | - | Furrow smoothing emissions factor | $E_{smooth}$ | kg CO<sub>2</sub> ha<sup>-1</sup> |
| Leaf area index | $LAI(t)$ | - | Turfing emissions factor (locally sourced) | $E_{turf,local}$ | kg CO<sub>2</sub> ha<sup>-1</sup> |
| Foliage biomass | $W_f(t)$ | t ha<sup>-1</sup> | Turfing emissions factor (imported) | $E_{turf,import}$ | kg CO<sub>2</sub> ha<sup>-1</sup> |
| Specific leaf area | $S_{LA}$ | m<sup>2</sup> kg C<sup>-1</sup> | Fertilisation emissions factor | $E_{fert}$ | kg CO<sub>2</sub> ha<sup>-1</sup> |
| Timestep | $n_{step}$ | Units time | Carbon stock (tissue/product X) | $S_X(t)$ | tCO<sub>2</sub> |
| Proportion of carbon allocated to foliage | $p_{foliage}$ | - | Initial carbon stock | $S_{X0}$ | tCO<sub>2</sub> |
| Leaf longevity | $l_{foliage}$ | yr | Carbon loss to atmosphere | $L_X(t)$ | tCO<sub>2</sub> |
| Gross primary productivity | $GPP(t)$ | C ha<sup>-1</sup> yr<sup>-1</sup> | Decay rate | $\alpha_X$ | yr<sup>-1</sup> |
| Net primary productivity | $NPP(t)$ | C ha<sup>-1</sup> yr<sup>-1</sup> | Half-life | $\lambda_X$ | Yr |
| NPP:GPP ratio | $Y$ | - | Decay efficiency | $\delta_X$ | - |
| Maximum light use efficiency | $\varepsilon$ | kg C MJ<sup>-1</sup> | Proportion of above-ground biomass allocated to product $X$ at harvesting time | $\rho_X(t_0)$ | - |
| Age dependent growth modifier | $f_{age}(t)$ | - | Total forest biomass at harvesting time | $C_{tot}(t_0)$ | tC |
| Annual average environmental growth modifier | $f_E$ | - | Forest biomass stored in tissue type or product $X$ at harvesting time | $C_X(t_0)$ | tC |
| Age at which $f_{age}(t) = 0.5$ | $A_{half}$ | yr | Power value of harvested biomass as biofuel | $\varepsilon_B$ | MWh t(Biomass)<sup>-1</sup> |
| **Water table / rooting depth modelling** | | | Grid-mix emissions factor | $E_{grid}$ | t CO<sub>2</sub> MWh<sup>-1</sup> |
| Forest soils emissions | $L_{forest}(t)$ | tCO<sub>2</sub> | Carbon : biomass ratio | $\rho_{C:B}$ | - |
| Unbounded rooting depth | $d_r'(t)$ | m | Biofuel plant efficiency | $\varepsilon_p$ | - |
| Rooting volume | $V_r(t)$ | m<sup>3</sup> | **Peatland restoration dynamics** | | |
| Root biomass | $B_r(t)$ | t(Biomass) ha<sup>-1</sup> | Restoration time | $t_{rest}$ | yr |
| Root biomass in 1 m<sup>3</sup> of soil | $\rho_{r,soil}$ | t(Biomass) m<sup>-3</sup> (soil) | Restoration shape parameter | $n_{rest}$ | - |
| Drainage depth | $d_d$ | m | **LCA summary statistics** | | |
| Peat depth | $d_p$ | m | Carbon payback time | $t_{payback}$ | yr |
| **Aquatic carbon losses** | | | Carbon flux intercept | $t_{flux}$ | yr |
| Total aquatic carbon loss rate | $L_{AqC}(t)$ | t CO<sub>2</sub> eq. ha<sup>-1</sup> yr<sup>-1</sup> | | | |

### B. 3PG parameterisation procedure

The parameterisation of the 3PG module used to represent lost sequestration potential is a crucial component of the LCA and therefore must be done carefully. The goal of the parameterisation procedure is to allow the user to input a value for the Yield Class (YC) of the forestry, or a reasonable proxy (e.g. height and age), and from this input, set the value of the time invariant global environmental modifier *f*_*E*_.

The first step was to run a battery of ‘full’ 3PG-SoNWaL simulations using the most up to date model dynamics (Robinson and O’Sullivan 2025, suzanneRobinsonFR/3PG-SoNWaL_FR) and species parameter estimates. 3PG-SoNWaL parameters were estimated by calibration against data from Alice Holt and from Gisburn 1 sample plots for Sitka spruce for Scots pine respectively. The Markov Chain Monte Carlo method (Ter Braak & Vrugt, 2008; Forrester et al. 2021) was used to estimate the free model parameters, and the 100 most likely parameter vectors were considered equally likely replicates for downstream analysis. For each of the 100 parameter vectors, the full model was run with the suite of environmental growth modifiers included in the full 3PG model (Landsberg and Waring, 1997; Xenakis et al. 2008; Almeida and Sands, 2016) fixed such that their product was equal to *f*_*E*_^∗^. Hereafter, *f*_*E*_^∗^denotes the global environmental modifier for the full 3PG-SoNWaL model, while *f*_*E*_is the equivalent parameter for the simplified model. Unfortunately, the two input parameters need to be considered independently because a simplified 3PG module parameterised by *f*_*E*_^∗^is typically more productive than a full model parameterised identically due to the absence of internal feedbacks between processes in the simplified model and the choice to ignore environmental fluctuations.

The full 3PG model tracks all relevant biomass compartments (stem, root and foliage) and uses allometric relationships to predict mensuration data for the stand (tree height, volume etc.). Therefore, for the full model, it is possible to assess YC according to the standard approach (Edwards & Christie, 1981): YC is defined by the annual stand volumetric increment (m^3^ ha^-1^ yr^-1^) at which the mean annual increment (MAI) and current annual increment (CAI) curves intersect. For simulations, LOESS regressions of volumetric increments against stand age were used to predict the average MAI and CAI curves across all 100 replicate parameter lists and their intersection assessed by geometric means (Fig B1, left). Since, MAI and CAI cannot be estimated for the simplified 3PG module which does not predict average tree dimensions, the peak in the NPP against time curve, available using both models, was also assessed by reference to the corresponding LOESS fit (Fig B1, right).

**Fig B1.**
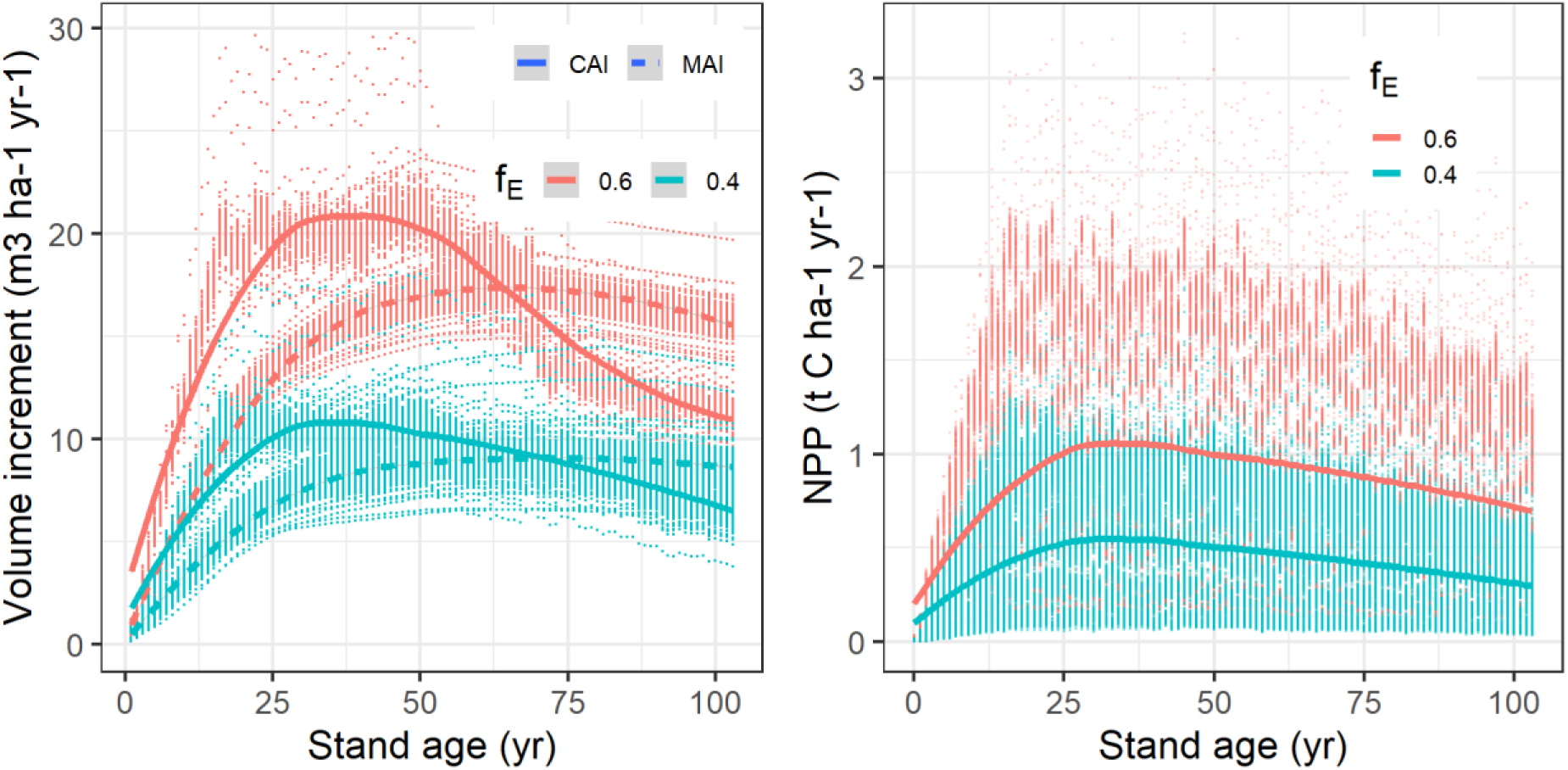
Yield class is estimated from the intersection of the LOESS regressions of Current Annual Increment (CAI) and Mean Annual Increment (MAI) in stand volume against stand age (left). Maximum NPP is estimated as the peak of the LOESS regression of NPP against stand age (right). Curves represent LOESS fits, points represent simulated values from each of the 100 replicate model runs. Note that NPP is a much more variable model output than volume increment since it is far more sensitive to environmental fluctuations in the full 3PG model.

With the YC estimated, a set of polynomial regressions were used to estimate an appropriate value of *f*_*E*_ for the user input YC. The required regressions were:

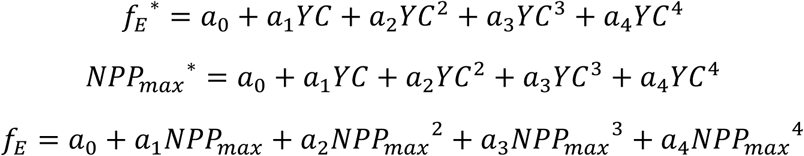

Here *NPP*_*max*_^∗^is the peak NPP for the full 3PG-SoNWaL simulation, while *NPP*_*max*_is the peak of the simplified model. Graphical representations of these polynomial associations are shown in Fig. B2. The estimated coefficients are shown in Tab. B1. Once the coefficients of these polynomial regressions were estimated, it was possible to predict a) *NPP*_*max*_^∗^, the peak NPP associated with the user input YC b) the value of *f*_*E*_ that produces the same peak, *NPP*_*max*_in the simplified 3PG model.

**Fig. B2.**
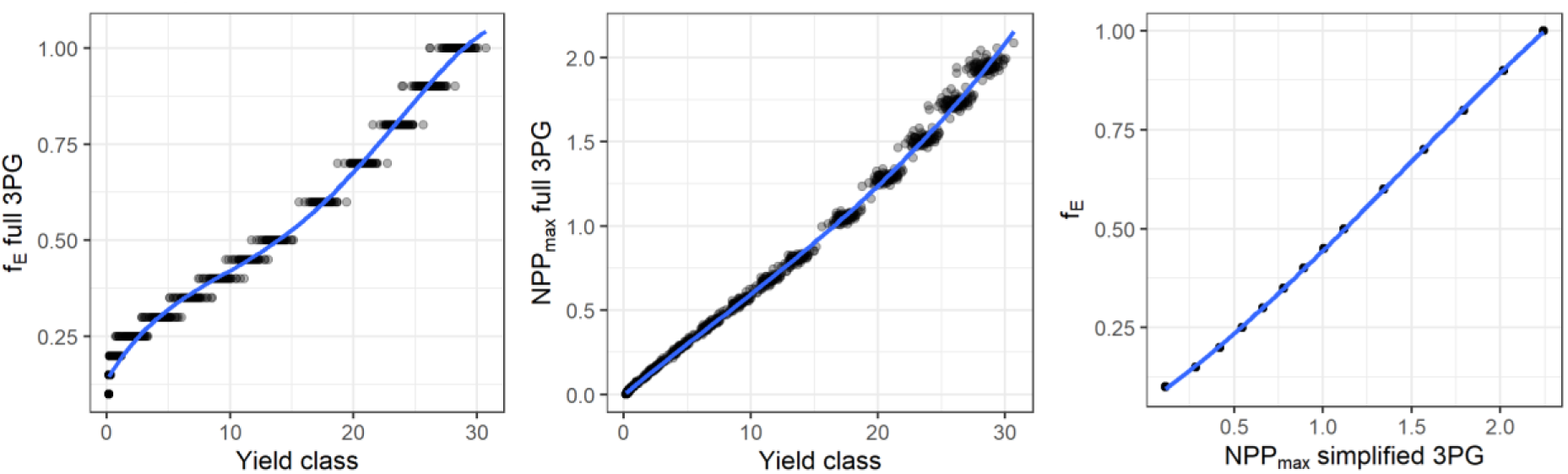
Emergent relationships between *f*_*E*_^∗^, *NPP*_*max*_^∗^and Yield class (YC) for the full 3PG model and between *f*_*E*_ and *NPP*_*max*_for the simplified 3PG model. The coefficients of the polynomial regressions shown in blue are used to set *f*_*E*_for the LCA based on user input YC.

**Tab. B1.** Polynomial regression coefficients estimated from analysis of the full and simplified 3PG models under variation of *f*_*E*_ ^∗^ and *f*_*E*_.

| $y$ | $x$ | Species | $a_0$ | $a_1$ | $a_2$ | $a_3$ | $a_4$ |
| --- | --- | --- | --- | --- | --- | --- | --- |
| $f_E^*$ | YC | S. spruce | 1.26E-01 | 7.91E-02 | -1.22E-02 | 9.89E-04 | -2.75E-05 |
| $NPP_{max}^*$ | YC | S. spruce | -1.85E-03 | 6.27E-02 | -5.52E-04 | 2.61E-05 | 0 |
| $f_E$ | $NPP_{max}$ | S. spruce | 7.22E-02 | 2.12E-01 | 2.64E-01 | -9.97E-02 | 0 |
| $f_E^*$ | YC | S. pine | 1.13E-01 | 1.38E-01 | -2.41E-02 | 6.55E-02 | -7.73E-05 |
| $NPP_{max}^*$ | YC | S. pine | 6.55E-02 | 8.45E-02 | 9.75E-03 | -4.45E-04 | 0 |
| $f_E$ | $NPP_{max}$ | S. pine | 1.25E-01 | 3.86E-01 | -7.25E-03 | 1.35E-02 | 0 |

### C. Forest growth modelling and emissions factors estimation

Yield tables for forest growth were based on ForestYield (Matthews et al., 2016). Wood product allocations from raw wood products to finished wood products (fuel, paper, wood-based panel, or sawn wood) was based on statistics from commercial forestry (Matthews et al., 2022). Proportions of biomass in crown, foliage and roots was based on expansion factors given in the CARBINE technical guide (Forest Research, 2025).

Emission factors for activities such as mulching, bund creation and ground smoothing were derived from existing sources.

#### Mulching

Forestry mulchers are universal brush cutters for cleaning forest areas from undesirable tree and shrub vegetation, for comminuting wood residues from logging, for cleaning fire lines, electrical overhead and underground cables (Marinov et al., 2023). From a review of existing mulchers on the market^3^, a number of them are based on mulching heads that are mounted on a tracked excavator or tractor (e.g Rabauld mulching heads^4^) which usually require an adaptable tractor of 80-300 hp (60-224 kW) for low-medium duty forestry operations.

Due to a lack of data on mulchers operating at biomass yields similar to that modelled in this study, fuel consumption was estimated based on work rates and fuel consumption of typical mulchers available on the market, though it was not possible to find both a work rate and fuel consumption rate for the same harvester.

Two values for work rate were found: 0.75 to 1.5 ha per 8 hour shift, or 0.1 to 0.2 ha/h for Prinoth Raptor 500^5^ (400 hp), and 0.57 acres/hour for a Caterpillar model 322B (161 hp, Halbrook et al. 2006), equivalent to 0.23 ha/h. Typical fuel rates of 26-34 litres/hour were based on two machines, the Prime Tech PT-300^6^ and the Fecon FTX300 mulching tractor^7^. Based on this, a fuel consumption rate of around 113 – 340 litres/ha or 303-911 kg CO2 ha-1- i.e. highly variable. This is based on the fastest work rate and lowest fuel consumption, and slowest fuel rate and highest fuel consumption.

#### Bund creation

The Peatland Action Technical Compendium (NatureScot, 2022) lists several stages of peatland restoration required to restore the hydrology of a peatland affected by drainage and/or alteration to its vegetation and wider surface integrity. These steps include:

- Building artificial dams or gullies to slowing the flow of water is often an initial necessity when restoring bare peat.
- Bunding (surface or below ground) is required to stop subsurface flow and water loss from an area or to stop surface flow of water to reduce water loss and erosion.
- Stabilisation and revegetation to stop further loss of peat from the bare peat surface (whether vertical or horizontal) by the combined actions of wind, water and oxidation, through re-establishing vegetation on such surfaces.

Emission factors for low carbon intensive options for rewetting and revegetating were derived from recent literature (Brennand et al., 2025), which demonstrated that the choice of intervention, source of materials, and the transport mechanisms used to deliver restoration can result in carbon costs that may be orders of magnitude different. Low carbon options include using peat dams and peat bunds to block drainage gullies; other higher carbon options can include building stone or plastic dams. Stone dams with materials delivered by helicopter are often recommended for what are claimed as remote and inaccessible locations, however the average emission factor per dam is estimated to be 135.94 kg CO_2_ eq, compared to 0.49 kg CO2 eq./ha for peat dams. The emission factor for plastic dams show the greatest variance in carbon costs due to several different plastic sheet options and the use of recycled PVC or new materials, ranging from 20 to 130 kg/dam. Emission factors per ha were based on a low intensive restoration plan as outlined by Brennand et al., (2025), requiring 65 dams and 2625 bunds for a 35-hectare site.

Revegetation options include heather brash spreading, and the spreading of imported or local turfs, with additional fertiliser and lime spreading. Higher carbon options, such as plug plants and the use of geojute were not considered. Geojute has shown promise on particularly sloped sites, where it can help hold bare peat in place so that it can be revegetated (Anderson, 2014).

**Tab. C1.** Emission factors for peatland restoration interventions in a low-carbon restoration model (Brennand et al., 2025).

|  |  |  | **Emission factor (kg CO<sub>2</sub> eq. ha<sup>-1</sup>)**<br>**(kg CO<sub>2</sub> eq. per intervention)** |  |  |
| --- | --- | --- | --- | --- | --- |
| **Stage** | **Technique** | **Intervention** | **Low** | **Medium** | **High** |
| Rewet | Gully/grip (drain) blocking | Peat dams | 0.46<br>(0.25) | 0.91<br>(0.49) | 1.86<br>(1) |
|  | Surface/shallow bunding | Peat bunds | 18.75<br>(0.25) | 25.5<br>(0.34) | 75<br>(1) |
| Revegetate | Direct establishment | Imported turf | 1516.42 | 1516.42 | 1516.42 |
|  | Direct establishment | Local turf | 471.76 | 471.76 | 471.76 |
|  | Indirect establishment | Lime and fertiliser | 1983.37 | 1983.37 | 1983.37 |

##### Ground smoothing

There are no studies giving the fuel consumption of stump flipping and ground smoothing for peatland restoration, however a study examining the time required for stump removal for bioenergy use were found (Laitila et al., 2008). From this, a typical work rate and fuel consumption can be derived for the initial stages of the process, including moving towards the stumps, positioning the boom to the stumps, lifting the stump and further processing and removing stumps. A work rate of 3.24 hours ha^-1^ of excavator time was estimated for site preparation following stump removal, which included mounding, compacting, and smoothing over of stump extraction holes. This represents around 20% of the time spent on the full process (therefore estimated to be 16.2 hours). Approximately another 30% of the total time is estimated to be required for moving and lifting the stumps. Therefore, a total time estimate for extracting stumps, flipping them and smoothing the surface would be approximately 8.1 hours ha^-1^.

A tracked excavator, typical for this operation, could be estimated to have a fuel consumption rate of 17.2-18.3 litres h^-1^ during a clearcut operation (Bertone & Mazone, 2025), meaning that the stump flipping and ground smoothing process could be estimated to consume around 139.3 and 148.2 litres fuel ha^-1^.

### D. Parameters used in the example LCA implementation

**Tab. D1.** Key inputs used in the scenario modelling shown in the main text.

| User input | Area 1 | Area 2 |
| --- | --- | --- |
| Management strategy | Harvested<br>Timber removed<br>Drainage dammed<br>Bunding installed<br>Furrows smoothed | Harvested<br>Timber left un-mulched on site<br>Drainage dammed<br>Bunding installed<br>Furrows smoothed |
| Area felled/restored (ha) | 1 |  |
| Tree species | Sitka spruce |  |
| Harvesting age (yr) | 50 |  |
| Yield class | Min: 6, Exp: 10, Max: 14 |  |
| Drainage depth (m) | 0.4 |  |
| Distance to processing sites (km) | 50 | - |
| Peat depth (m) | 2 |  |
| Fallow period between harvesting and restoration (yr) | 5 |  |
| Peatland ecosystem function restoration time (yr) | 50 | 100 |
| Peatland ecosystem function restoration shape parameter | 1.5 | 4 |
| Pre-restoration water table depth (m) | 0.5 |  |
| Post-restoration water table depth (m) | Min: 0.01, Exp: 0.05, Max: 0.10 |  |

### E. Detailed sensitivity analysis

**Fig. E1.**
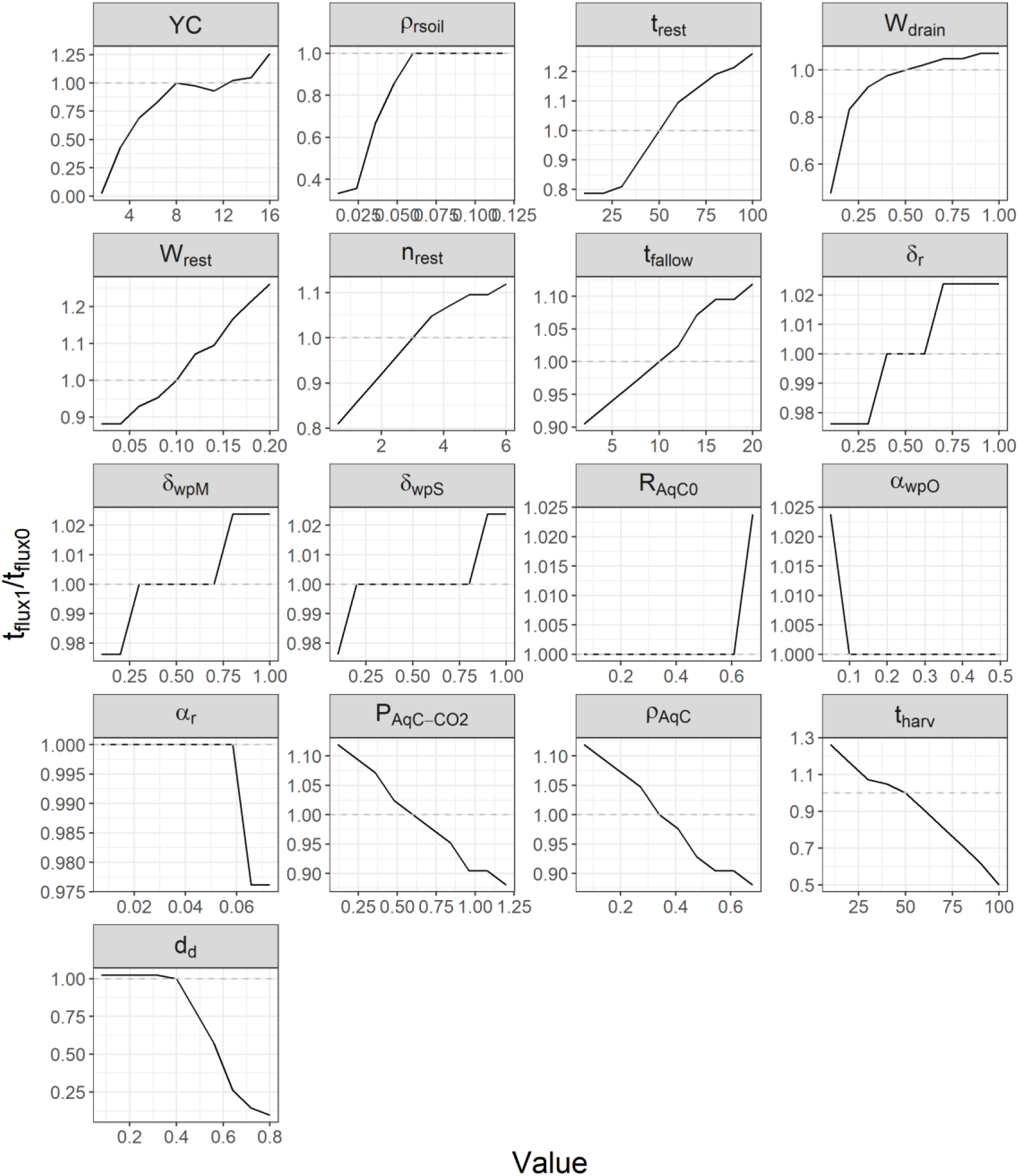
Relative change in the carbon flux intercept under variation of input variables. Only variables whose variation produces change of at least 1 year are shown. Step changes in mitigation time are due to the minimum temporal resolution of 1 year.

**Fig. E2.**
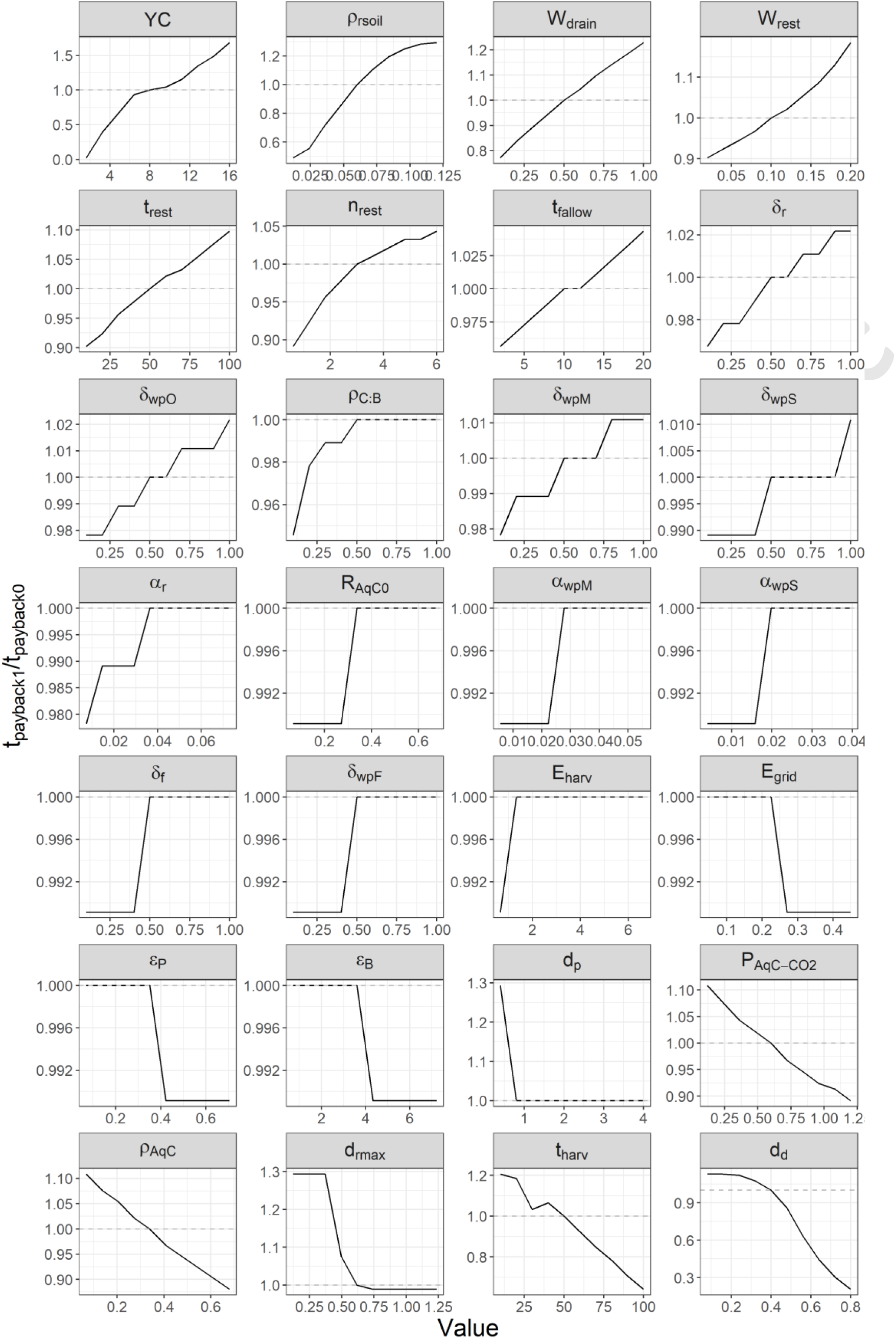
Relative change in the carbon payback time under variation of input variables. Only variables whose variation produces change of at least 1 year are shown.

